# Distinct Roles of Deep and Superficial Cortical Layers in Tone Prediction, Comparison, and Adaptation in Human Auditory Cortex

**DOI:** 10.1101/2025.11.24.687809

**Authors:** Jorie van Haren, Floris P. de Lange, Sonja A. Kotz, Federico De Martino

## Abstract

Auditory perception relies on prior sensations both to form expectations about upcoming sounds and to adapt to repeated inputs. Yet, how these processes interact with bottom-up frequency-specific stimulus drive, and how they are organised across cortical depth, remains unclear. Using ultra-high-field functional magnetic resonance imaging (7T fMRI), participants listened to stochastic tone sequences whose repetition structure and predictability varied over time. For each voxel independently, we used computational modelling to estimate responses for stimulus drive, repetition suppression, and expectations. We then partitioned the unique and shared variance of these components to estimate their relative importance in modulating BOLD activity across cortical layers in auditory cortices. Expectation-related signals explained most unique variance in deep (infagranular) layers, in line with accounts that place internal model representations and belief updates in deeper cortical populations. Variance jointly explained by stimulus drive and expectation was instead strongest in superficial layers, consistent with these compartments expressing computations that combine bottom-up input with top-down predictions, such as prediction error signalling. Finally, repetition suppression accounted for variance uniformly across depth, suggesting an adaptation-like mechanism that modulates responsiveness without a clear laminar bias. Together, the data suggest that deep layers contain content-specific predictive models, superficial layers register prediction–input alignment, and repetition suppression provides a depth-invariant, local gain control that complements predictive processing.

## 1 Introduction

Sound perception takes place in ever-changing acoustic environments, where the brain must piece together information that occurs over different time periods [1, 2]. Contextual cues from both the recent and distal past are used to generate predictions about what is likely to occur next. As new evidence arrives, these predictions are updated, yielding a continuously evolving internal model of the auditory environment [3, 4]. Alongside these forward-looking processes, simpler local mechanisms such as stimulus-specific adaptation attenuate responses based solely on recent acoustic history [5, 6, 7, 8]. Because predictions frequently coincide with the repetition of low-level features, adaptation-driven attenuation and prediction often co-occur, making it difficult to disentangle repetition suppression from prediction-related modulation at the neural level [7, 9].

To understand how these processes are implemented in the human brain, we must consider the mesoscopic circuitry through which stimulus-specific adaptation and forward-looking predictive signals influence neural activity. Feedback projections from higher-order regions primarily target deep layers of sensory cortices [10, 11, 12]. These projections carry predictions that reflect content-specific organization, such as anticipated frequency patterns [13, 14, 10]. In turn, feedforward pathways relay prediction errors from superficial layers upward to support model updating [15, 16]. Repetition suppression is often proposed to arise locally, reflecting a bottom-up mechanism that selectively dampens responses within frequency-tuned populations [17, 18], yet has also been shown to vary with predictive context [19]. This laminar organisation suggests that predictive signals and repetition suppression may not only differ in their cortical depth profiles, but also in how they act on specific stimulus content.

Content specificity in the processing of acoustic frequencies can be related to the tonotopic organisation of auditory cortices, where both adaptation and predictive processes are anchored to acoustic frequency bands [20, 21, 22]. When a particular frequency recurs, repetition suppression attenuates the responses of the neuronal populations tuned to that frequency [2, 5, 6]. At the same time, predictions about upcoming frequency ranges can selectively modulate these same populations, shaping their responses even before the stimulus appears [11, 12]. Moreover, rather than settling on a single prediction, the brain likely maintains a probabilistic model with overlapping expectations over multiple outcomes [23, 24, 25, 9]. Tracking these distributed predictions ensures that cortical activity reflects not just the most likely event but also a spectrum of alternative predictions [23, 24, 25, 9]. Probing frequency-tuned responses under stochastic conditions makes it possible to move beyond static receptive-field descriptions and quantify how stimulus drive, repetition suppression, and top-down predictions jointly shape cortical responses and perception.

Traditional oddball paradigms have provided valuable insights into how auditory mismatch responses can reflect both adaptation and prediction [26]. The discrete violations used in these paradigms, however, capture only part of the processes engaged during natural listening, where statistical regularities evolve dynamically rather than occurring as isolated events [27]. In such contexts, the brain must maintain and continuously update internal models to track shifting regularities, adjusting predictions based on both transient deviations and broader structural changes [1, 28, 29]. This ongoing recalibration highlights the need for approaches that move beyond static contrasts, capturing how predictive signals and adaptation mechanisms jointly evolve as the acoustic context changes. Under these conditions, adaptation and prediction periodically diverge, offering a window into how their distinct contributions unfold in time and across cortical depth.

To understand how the brain maintains accurate models of our dynamic sensory environment, it is necessary to establish how adaptive and predictive mechanisms each contribute to acoustic processing within the tonotopic landscape and mesoscopic circuitry [18, 30]. To quantify their proportional contributions, we leveraged temporally unstable stochastic tone sequences. Specifically, we created a dynamic auditory environment in which tones were sampled from two Gaussian distributions whose sampling probabilities shifted over time. This design produced sequences that alternated between stable periods — eliciting strong repetition suppression — and transitions that required listeners to update their internal models [9, 28]. To capture these processes quantitatively, we employ a computational modeling framework that dynamically tracks the moment-to-moment predictive probability estimates of each sound-related frequency and integrates tonotopic representations of repetition suppression driven by recent history [6, 8], current stimulus drive [21], and future-oriented inference [31]. We then used variance partitioning to estimate how much of the cross-validated variance in cortical activity each mechanism uniquely accounted for, and how much was shared — revealing their relative importance in shaping the measured BOLD responses [32]. Crucially, the shared components highlight how predictive inference and repetition suppression intersect with frequency-driven input responses, offering a window into the integration of top-down and bottom-up processes across cortical depth. By combining these model-derived estimates with submillimeter-resolution fMRI, we examined how predictive inference and repetition suppression jointly shape cortical responses across layers. This approach reveals how local repetition suppression complement top-down predictions, enhancing sensitivity to broader statistical changes and improving the brain’s ability to anticipate future events.

## 2 Results

Ten participants were recorded using ultra-high-field functional magnetic resonance imaging (fMRI) - at 7 Tesla - over two separate sessions. In the first session, participants listened to auditory sequences of pure tones drawn from two Gaussian distributions with shifting sampling probabilities (two sessions each; see Figure 1d). We investigated how these dynamic auditory sequences recruit stimulus drive, repetition suppression, and expectation mechanisms across cortical layers in the auditory cortices in a content-specific, tuning-dependent manner. To establish the frequency-selective tuning properties independently of contextual influences from repetition suppression and expectation, we used quasi-random tone sequences as a functional localizer for population receptive field (pRF) mapping (session 1: see Figure 1a). Localizer sequences spanned approximately six octaves and were designed to avoid short-term repetitions and statistical structure. Functional pRF mapping results, aligned using curvature-based registration of individual cortical surfaces, are presented as group averages for both hemispheres in Figure 1b. Additionally, pRF tuning width measurements are shown in the same Figure 1c. Individual functional pRF estimates for all ten participants are available in Supplementary Figure **S4**. The resulting tonotopic maps reveal clear, mirror-symmetric frequency gradients centred on the crown of Heschl’s gyrus (HG) or within the sulcus intermedius [33].

**Figure 1.**
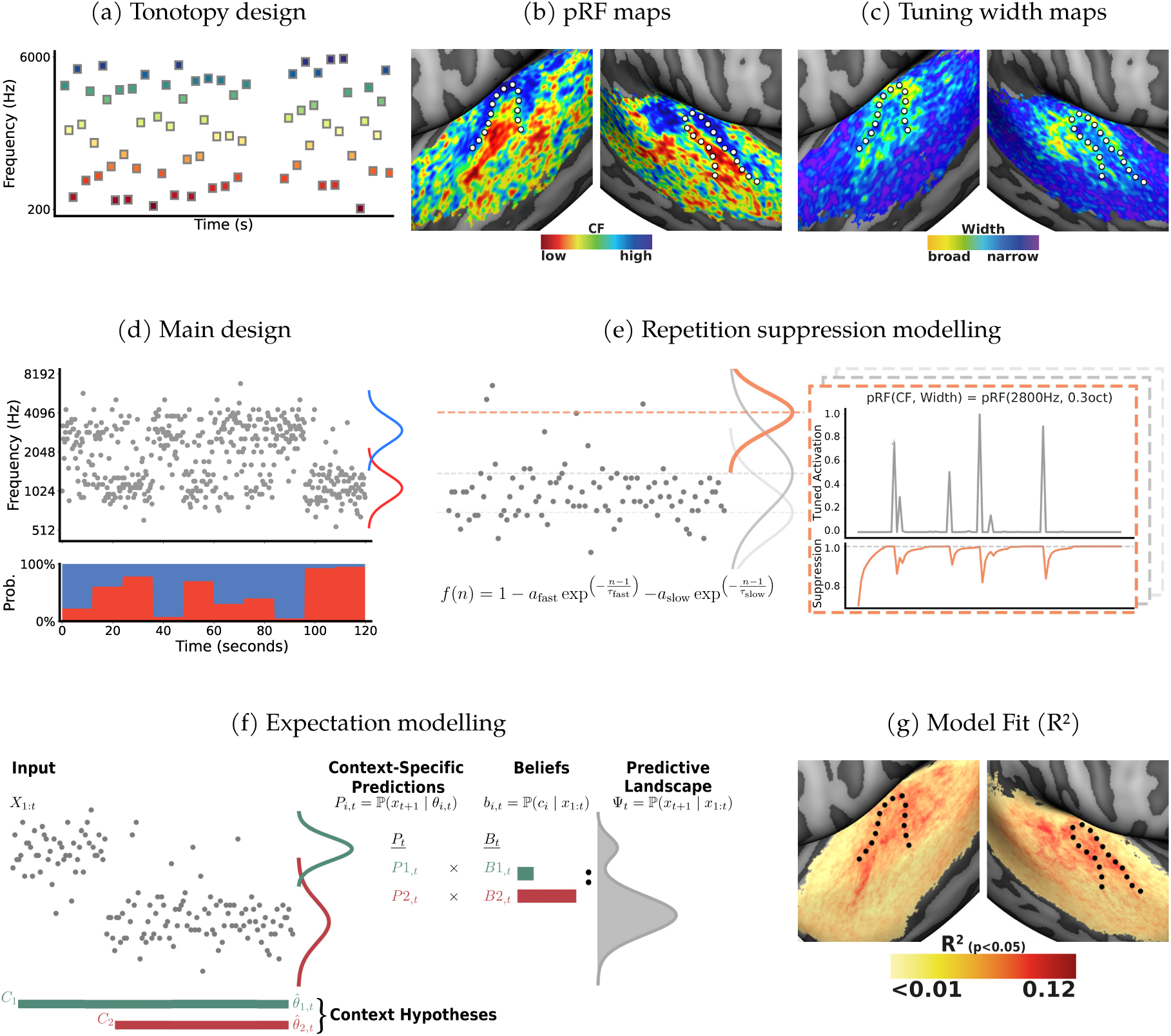
Quantifying Tuning-Based Responses to Dissociate Repetition Suppression and Predictive Inference. **a** Tonotopy localizer: amplitude-modulated pure tones (1.4 s duration, 0.4 s inter-stimulus interval) were presented in quasi-random order, minimising short-term repetition and statistical structure to estimate frequency-selective population receptive fields (pRFs). **b** Group-average pRF maps, masked to auditory cortex and thresholded at *r* > .2 based on the correlation between Gaussian pRF model predictions and voxelwise single beta estimates. The dotted white line denotes the anatomical border of Heschl’s gyrus. **c** Corresponding tuning width maps derived from pRF fits. **d** Main paradigm: pure tones (200 ms duration, 50 ms inter-stimulus interval) were sampled from mixtures of two Gaussian distributions separated by 1.48 octaves. The upper panel shows the presented tone frequencies; the lower panel shows the underlying sampling probabilities. **e** Repetition suppression modeling: a long-trace adaptation model estimated voxelwise responses based on recent stimulus history, weighted by prior activation within each voxel’s frequency-selective profile. Suppression followed a double-exponential decay across trials, capturing both rapid and sustained adaptation. These history-dependent weights modulated each voxel’s response over time, producing time-resolved predictions of frequency-specific suppression. Grey and orange Gaussians illustrate example voxel tuning functions (pRFs): orange corresponds to the inset example (pRF(2800 Hz, 0.3 oct)). **f** Expectation modeling: the D-REX model [31] computed a dynamic predictive landscape (Ψ_*t*_) by integrating predictions across multiple context hypotheses. Each hypothesis was defined by local summary statistics *θ*_*i,t*_ (e.g., mean and variance) and generated a context-specific prediction *P*_*i,t*_ = ℙ (*x*_*t*+1_ | *θ*_*i,t*_), weighted by its belief *b*_*i,t*_ = ℙ (*c*_*i*_ *x*_1:*t*_). The resulting predictive landscape was used to derive tone-by-tone surprisal, precision, and prior expectations across the frequency space. **g** Model fit (*Stimulus Drive ⋃ Repetition Suppression ⋃ Expectations*): group-averaged *R*^2^ maps showing the variance explained by the full model across left and right hemispheres. Maps were manually masked to auditory cortex and thresholded at *p* < .05 (uncorrected). The dotted black line denotes the anatomical border of Heschl’s gyrus. Colour indicates model fit, with red reflecting higher *R*^2^ values and yellow lower values (range *≈* 0–0.12).

The population receptive field (pRF) estimates of each voxel’s preferred centre frequency and tuning width were used as inputs for computational modeling of stimulus drive, repetition suppression, and expectation-driven activity. These models quantified, for each voxel separately, how strongly it was expected to respond to each tone in the main experimental runs based on its population frequency tuned stimulus drive, recent stimulation history (repetition-suppression), or evolving predictions about upcoming tones (session 2: see Figure 1e & 1f). The resulting predicted time courses were then entered as regressors in cross-validated linear regression analyses, allowing to estimate how much of the observed BOLD signal could be explained by each of the three processes. To disentangle their relative contributions, we applied a set-theoretic variance partitioning procedure [32], quantifying unique and shared variance attributable to each component and their combinations. Through this framework, we could assess how much variance each process explained above and beyond the others, providing a quantitative account of their relative contributions to auditory cortical responses.

The joint model, incorporating all three components, accounted for a significant proportion of variance across bilateral temporal cortices (see figure 1g), with cross-validated *R*^2^ values in held-out data reaching up to approximately 12%, confirming it captured meaningful aspects of the responses. Partitioning this variance into its component processes revealed robust and widespread contributions of both stimulus drive (≈ 5–20%of the cumulative explained variance (EV); *p*_FDR_ ≤ 2.4e−2) and repetition suppression (≈ 9–17%EV; *p*_FDR_ ≤1.7e−4) across auditory cortices, with expectation effects emerging more selectively toward higher-order regions (≈ 3–23%EV; *p*_FDR_ ≤ 8.3e−3). Detailed region-wise results are provided in Supplementary analysisA.1.

### 2.1 Distinct Laminar Profiles of Expectations Trending Toward Deep Layers and Layer-Uniform Repetition Suppression

Having established robust contributions of stimulus drive, repetition suppression, and expectation across the auditory cortices, we next examined whether these processes exhibit distinct laminar profiles. To this end, we focused on five bilateral regions of interest (ROIs): Heschl’s gyrus (HG), Planum Polare (PP), Planum Temporale (PT), and the anterior and posterior Superior Temporal Gyrus (aSTG, pSTG). Each ROI was subdivided into deep, middle, and superficial compartments using an equi-volume approach [34].

The variance explained by stimulus drive showed no significant variation across cortical depth in Heschl’s gyrus (HG), Planum Polare (PP), or Planum Temporale (PT), indicating a uniform engagement of frequency-selective responses across primary and belt regions. The same held true for the anterior Superior Temporal Gyrus (aSTG), where unique variance explained by the stimulus drive was evenly distributed across layers. Only posterior STG (pSTG) showed a consistent depth-related modulation, with greater unique variance in deep compared to middle layers (*p*_FDR_ = 1.5e−3; FDR-corrected across 15 comparisons spanning 5 ROIs and 3 layer contrasts) (see figure 2). This pattern may reflect that frequency-specific representations are more distinct in deep layers, whereas superficial layers may support more integrated codes that combine frequency with additional spectrotemporal features [35] or more complex tuning [36, 37]. Overall, stimulus drive-related signals appeared broadly expressed across cortical depth, with only limited evidence for laminar specialization.

**Figure 2.**
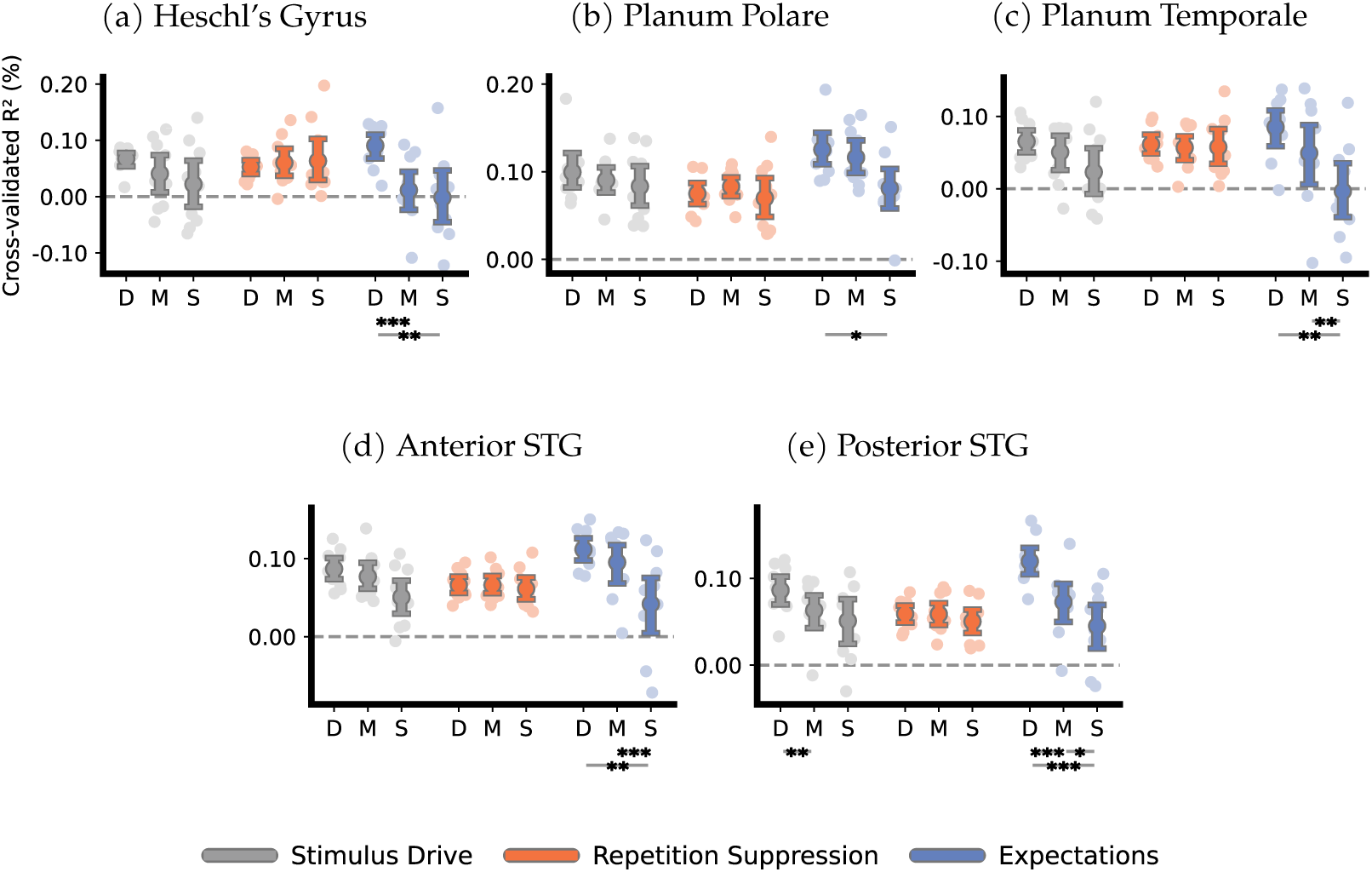
Laminar dissociation of stimulus drive, repetition suppression, and expectation: deep-layer preference for expectations across auditory cortex. Each panel shows the cross-validated unique variance (variance accounted for by each model beyond what could be explained by the other two) explained by the stimulus drive, repetition suppression, and expectation components across cortical layers. D, M, and S denote deep, middle, and superficial layers, respectively. Results are shown for five auditory regions of interest (ROIs): **a** Heschl’s Gyrus (HG), **b** Planum Polare (PP), **c** Planum Temporale (PT), **d** anterior Superior Temporal Gyrus (aSTG), and **e** posterior STG (pSTG). Stimulus drive and repetition suppression exhibited no consistent variation across cortical depth, whereas expectation-related activity showed a robust and selective deep-layer bias. Stars indicate significance levels of the cross-validated unique variance explained across depth (paired-sample bootstrap t-tests, FDR-corrected across all layer pair comparisons and ROIs): *p*_FDR_ *<* 0.05 (*), *p*_FDR_ *<* 0.01 (**), *p*_FDR_ *<* 0.001 (***). Error bars indicate bootstrapped 95% confidence intervals.

Repetition suppression similarly showed no substantial variation in its variance explained across cortical layers in HG, PP, PT, aSTG, and pSTG (see figure 2). This consistent pattern supports the view that repetition suppression reflects a locally implemented adaptation process, expressed evenly across cortical depth. A complementary analysis presented in the Supplementary Material (Section A.3) further characterizes the temporal profile of this adaptation process, revealing a progressive shortening of the repetition-suppression window from primary to associative auditory regions.

The portion of the signal explained by expectations, in contrast, displayed a robust and systematic laminar gradient (see figure 2). In HG, more unique variance was explained in deep compared to middle (*p*_FDR_ = 6.0e−4) and deep compared to superficial layers (7.2e−3), revealing a depth-specific organization of the modulation by expectation. A similar infragranular preference was present in PP (deep > superficial:*p*_FDR_ = 3.7e− 2) and PT (deep > superficial:*p*_FDR_ = 7.3e−3; middle > superficial: *p*_FDR_ = 7.4e−3). This laminar profile became even more pronounced in associative regions. In aSTG, expectation-related unique variance was greater in both deep and middle compared to superficial layers (*p*_FDR_ = 2.2e−3and6.0e− 4), likewise in pSTG, we observed a similarly consistent gradient, with more unique variance explained in deep versus superficial (*p*_FDR_ = 6.0e−4), deep versus middle (*p*_FDR_ = 6.4e−4), and middle versus superficial layers (*p*_FDR_ = 1.1e−2). This consistent gradient across primary, belt, and associative areas highlights a clear laminar organisation specific to expectation signals, distinguishing them from the more uniformly distributed stimulus drive and suppression components.

Given this consistent infragranular profile, a question is whether the observed deep-layer bias of expectation-related variance primarily reflects the priors represented within the expectation model. To address this, we conducted an additional analysis in which priors and prediction errors were modelled separately relative to a stimulus drive - repetition suppression baseline (see Supplementary analysisA.2). This adjustment necessarily came at the cost of the shared contribution components examined below. Both priors and prediction errors independently exhibited preferential expression in deep layers across the auditory hierarchy (minimum *p*_FDR_ ≤ 1.0e−5for errors; *p*_FDR_ ≤ 5.7e−3for priors), with priors showing a more region-specific pattern. Consistent with this, a coefficient-level analysis revealed no reliable laminar dissociation between priors and errors (Figure **S5**). These results indicate that the infragranular expectation effects reported here are not solely driven by top-down priors but likewise encompass prediction-error computations expressed within the same deep-layer circuitry, consistent with predictive-coding architectures in which deep populations maintain internal models of the sensory environment together with their associated errors [15, 38, 30, 10, 39].

Together, these findings reveal a clear functional dissociation across cortical depth. Expectation-related contributions consistently followed a depth-dependent gradient across primary, belt, and higher-order auditory regions, with greater unique variance explained in deeper compared to superficial layers throughout the auditory hierarchy. This laminar profile aligns with hierarchical predictive coding accounts, which propose that deep layers are the principal recipients of top-down predictions. In contrast, stimulus drive and repetition suppression exhibited relatively uniform expression across cortical depth, with no strong bias toward the middle layers typically associated with bottom-up input. These results suggest that, unlike expectation, stimulus-driven and adaptation processes reflect locally implemented computations that are distributed relatively evenly across cortical depth, without clear laminar specialization.

### 2.2 Superficial Layers Represent Shared Expectation–Stimulus Drive Dynamics

Having found that the expectation model uniquely explained the most variance in infragranular layers, we next asked whether combinations of computational processes are likewise organised with respect to cortical depth. Crucially, the shared components reveal how predictive inference and repetition suppression intersect with frequency-specific stimulus drive. These intersections capture portions of the signal where the modeled processes jointly account for fluctuations, isolating shared explanatory structure in the data that neither model alone, nor variance shared across all three components, can explain. To this end, we examined the variance jointly explained by each pair of computational models, excluding variance shared with all three. Specifically, we tested shared variance between the stimulus drive and repetition suppression models (*Stimulus Drive* ∩ *Repetition Suppression*), between the stimulus drive and expectation models (*Stimulus Drive* ∩ *Expectation*), and between the repetition suppression and expectation models (*Repetition Suppression* ∩ *Expectation*).

The variance shared between stimulus drive and expectation (*Stimulus Drive* ∩ *Expectation*) - reflecting variance that can only be explained by their joint contribution, and crucially not by repetition suppression alone - consistently exhibited a bias toward superficial layers across auditory cortices (see Figure 3). In Heschl’s gyrus (HG), superficial layers accounted for more shared variance than deep layers (*p*_FDR_ = 4.6e− 3; FDR-corrected across 15 comparisons spanning 5 ROIs and 3 layer contrasts). A similar, albeit weaker, pattern was present in Planum Polare (PP), where expectations and stimulus drive together tended to explain more variance toward the surface but did not reach significance after FDR correction (superficial > deep: *p*_FDR_ = 6.8e−2). In Planum Temporale (PT), a clear superficial bias again emerged, with greater jointly explained variance in superficial compared to both deep and middle layers (*p*_FDR_ = 1.3e−3for both). In anterior superior temporal gyrus (aSTG), the same pattern was evident (superficial > deep, *p*_FDR_ = 1.5e− 2). Finally, in posterior STG (pSTG), a graded pattern emerged across all layer pairs, with more shared variance in superficial compared to deep (*p*_FDR_ = 6.0e−4), superficial compared to middle (*p*_FDR_ = 5.1e−2), and middle compared to deep layers (*p*_FDR_ = 6.0e− 4). This superficial layer bias indicates that the joint influence of frequency-selective input drive and predictive expectations is more strongly expressed toward the cortical surface, in line with computations involved in prediction error calculation [15, 16].

**Figure 3.**
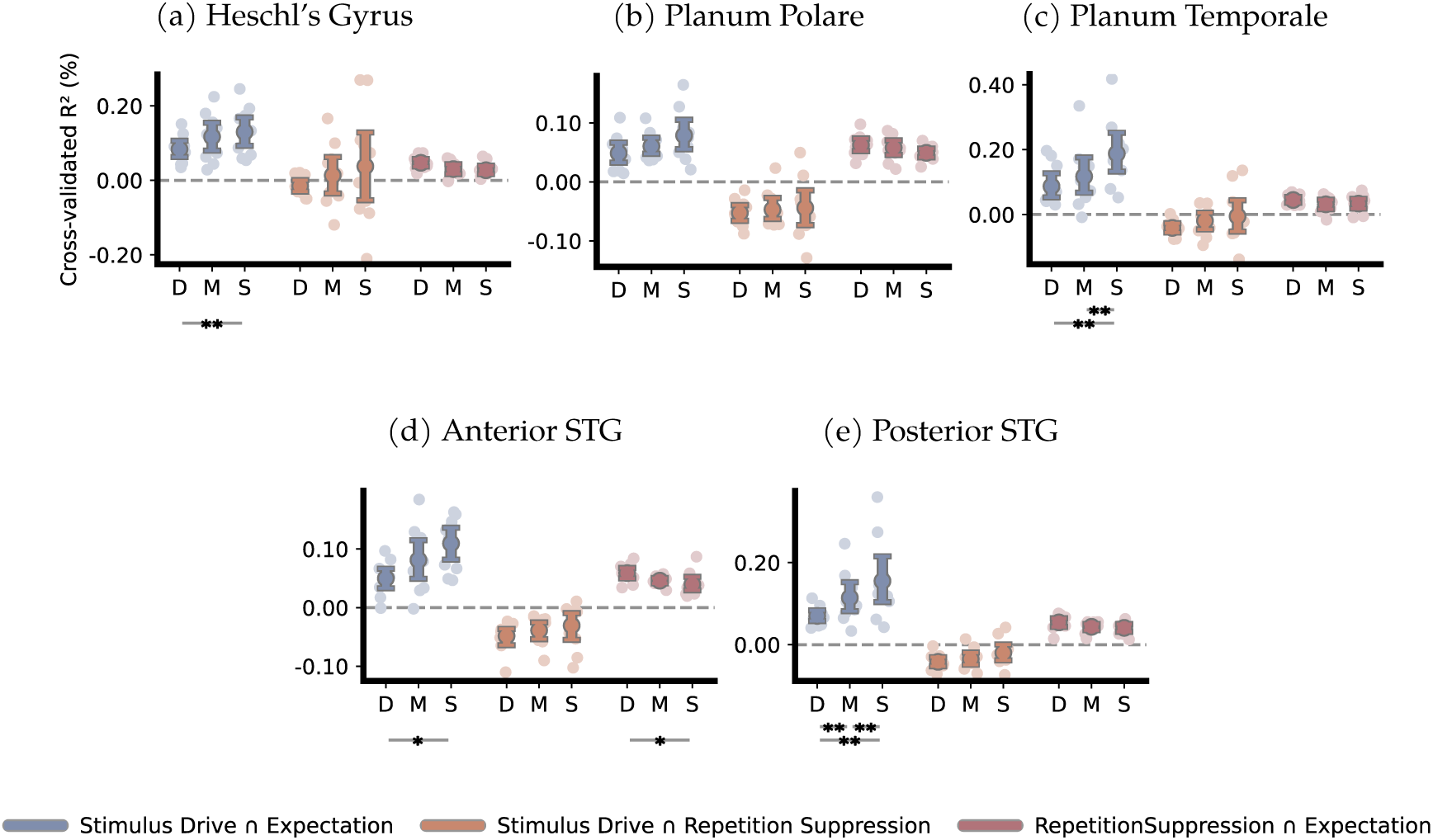
Laminar profiles of shared variance: widespread superficial stimulus drive–expectation bias and focal deep effects for repetition suppression–expectation. Each panel shows the cross-validated unique variance jointly explained by pairs of model components, excluding variance explained by the third, across cortical layers. D, M, and S denote deep, middle, and superficial layers, respectively. Shown are: *Stimulus Drive* ∩ *Expectation, Stimulus Drive* ∩ *Repetition Suppression*, and *Repetition Suppression* ∩ *Expectation*, computed via variance partitioning. Results are displayed for five auditory regions of interest (ROIs): **a** Heschl’s Gyrus (HG), **b** Planum Polare (PP), **c** Planum Temporale (PT), **d** anterior Superior Temporal Gyrus (aSTG), and **e** posterior STG (pSTG). *Stimulus* ∩ *Drive* ∩ *Expectation* exhibited a consistent superficial-layer bias across regions, whereas *Repetition Suppression Expectation* showed a localised deep-layer bias in aSTG. Shared variance between stimulus drive and repetition suppression showed no systematic laminar pattern. Stars indicate significance levels of the cross-validated unique shared variance across depth (paired-sample bootstrap t-tests, FDR-corrected across all layer pair comparisons and ROIs): *p*_FDR_ *<* 0.05 (*), *p*_FDR_ *<* 0.01 (**), *p*_FDR_ *<* 0.001 (***). Error bars indicate bootstrapped 95% confidence intervals.

In contrast, shared variance between repetition suppression and expectation (*Repetition Suppression* ∩ *Expectation*) was uniformly expressed across depth for most regions (see figure 3) However, in anterior STG (aSTG), it exhibited the opposite pattern to that we observed for *Stimulus Drive* ∩ *Expectation*, with more unique shared variance explained in deep compared to superficial layers (*p*_FDR_ = 2.1e−2). This localised infragranular bias suggests that when adaptation and prediction mechanisms converge, their intersection may preferentially engage deeper cortical circuits.

Shared variance between stimulus drive and repetition suppression (*Stimulus Drive* ∩ *Repetition Suppression*) did not vary systematically with depth in any region (see figure 3). Across all ROIs, this intersection term was uniformly expressed across cortical layers, indicating that the joint expression of stimulus drive and repetition suppression reflects a depth-invariant process.

Taken together, these results reveal distinct laminar profiles: when predictive information is integrated with frequency-selective sensory input, the resulting signal is preferentially expressed towards superficial layers across multiple auditory areas. In contrast, when prediction overlaps with repetition suppression, a significant depth effect — favouring deeper layers — is observed in aSTG, although this effect remains localised. These patterns align with hierarchical predictive coding theories, positing that superficial compartments preferentially encode prediction errors by integrating top-down expectations with bottom-up sensory evidence [15, 38, 30, 10, 39].

## 3 Discussion

We combined ultra-high-field fMRI with voxel-wise - frequency-specific - models of stimulus drive, repetition suppression, and expectation dynamics to characterise their relative contributions to responses evoked by dynamic tone sequences across cortical layers. This mapping revealed a consistent deep-layer bias for unique expectation signals across primary, belt, and associative auditory cortices, contrasted with depth-invariant profiles for stimulus drive and repetition suppression. At the same time, across these regions, variance jointly explained by stimulus drive and expectation was preferentially expressed in superficial layers, indicating that predictive modulation and stimulus drive converge most strongly near the cortical surface.

Repetition suppression consistently shaped responses across the auditory hierarchy, reflecting a robust sensitivity to recent acoustic history. This suppression closely mirrored stimulus drive and exhibited no clear laminar gradient, suggesting that it operates uniformly throughout depth and is largely independent of hierarchical feedback. In contrast, expectation-related signals were strongest in infragranular layers, signalling a more global process reliant on feedback from higher-order regions to integrate contextual information [10, 11, 12]. Collectively, within the soundscapes we investigated, the observed dynamics reflect a convergence of stimulus drive, local adaptation, and globally informed predictive inferences, combined within a tuning-dependent framework.

Across auditory cortices, expectation-related signals were consistently expressed towards deep cortical layers when considering their unique variance contribution. This laminar profile indicates that the internal model guiding auditory predictions is maintained within infragranular circuits. These findings align with canonical microcircuit accounts which assign deep-layer populations the role of representing contextual information and propagating prediction signals through descending pathways [15, 38, 30, 10, 39]. In this framework, the localization of expectation signals to deep layers reflects their role as a primary recipient site of top-down inferences containing integrated contextual structure over time. Together, these observations support hierarchical predictive-coding models, in which infragranular populations not only broadcast but also continually revise auditory predictions as sensory input unfolds [10, 40]. Crucially, these deep-layer signals extended beyond frequency-tuned acoustic responses, pointing to a model-based component of auditory inference not reducible to input-driven selectivity.

Although prediction errors are typically linked to superficial cortical layers [15, 30], we found no evidence for a distinct superficial component that could be attributed to prediction-error signals independent of stimulus-driven responses. Instead, expectation-related variance in superficial layers emerged only when it coincided with frequency-selective stimulus drive, suggesting that these superficial responses reflect the comparison between predicted and received input rather than a separable error signal. In other words, this variance was confined to populations engaged by the presented stimulus and did not manifest in predicted-but-unengaged populations, exemplifying the frequency-specific nature of error computations in predictive processing. Hence, while deep layers host top-down generated predictions and their updates independent of sensory input, superficial layers may serve as a locus where those predictions are tested against incoming stimuli. To further clarify the nature of this superficial-layer effect, paradigms that introduce mismatches without concurrent sensory drive - for example, through unpredictable omissions - may help determine whether superficial layers represent a dedicated error signal or merely instantiate the comparison process itself.

A central theme in our results is that all measured mechanisms — stimulus drive, repetition suppression, and predictions — are grounded in tuning, making their expression inherently content-specific. Rather than collapsing onto a single prediction or suppression trace, auditory cortices maintain a representational landscape in which tuned units carry both history-dependent suppression profiles and probabilistic expectation profiles. Repetition suppression leaves lasting shadows across multiple tones, shaping a frequency-anchored suppression landscape that operates alongside prior distributions. Our stochastic sequences demonstrate that perception is not governed by a single prediction, nor dampened within a single unit, but instead reflects the joint influence of tuning-anchored histories and probabilistic forecasts - a distinction that may be obscured in designs or analyses that average over tuned subpopulations. Overall, these mechanisms construct a structured representational landscape that flexibly combines recent input, long-lasting suppression, and probabilistic predictions. This landscape encodes graded likelihoods across frequency-selective populations, enabling perception to integrate recent history with contextual uncertainty.

Together, our results indicate that auditory cortices implement content-specific computations with distinct laminar footprints. Deep layers carry the internal model: a frequency-tuned predictive landscape of probabilistic predictions and their updates that bias processing across the hierarchy. Superficial layers, in contrast, preferentially express the alignment between content-specific expectation and sensory input, consistent with a comparison stage in which predicted and received evidence are jointly represented and expressed when sensory drive is present. Alongside these predictive signals, repetition suppression acts as a pervasive, locally implemented and laminar-uniform mechanism that modulates gain based on recent history without engaging hierarchical message passing. This division of labour suggests a compact scheme for inference in dynamic soundscapes: deep, content-specific predictions continuously shape the operating point of the system; superficial circuits register their consequences at stimulus arrival; and locally uniform adaptation compresses recent context. In combination, these mechanisms allow the auditory cortex to remain sensitive to broad statistical changes while efficiently tracking immediate repetitions. These parallel, content-specific processes modulate tuned responses across auditory cortex — allowing predictions and local adaptation to selectively target distinct neural populations, with repetition suppression providing a fast, efficient modulation that complements more complex predictive inferences.

## 4 Methods

### 4.1 Participants

Ten participants (five female; *M* = 22.6 years, age range: 18–29 years) participated in this fMRI experiment. They reported normal hearing and no history of neurological disorders. The study was approved by the Ethics Review Committee of the Psychology and Neuroscience faculty (ERCPN; Approval Code: OZL-232-01-01-2021) at Maastricht University and was conducted in accordance with the principles of the Declaration of Helsinki. Written informed consent was obtained from all participants prior to the start of the experiment.

### 4.2 MRI Acquisition

MRI data were acquired on a 7-Tesla Siemens MAGNETOM scanner using a 32-channel head coil (Nova Medical) at Scannexus (Maastricht, The Netherlands).

Session 1: High-resolution anatomical data were collected, including a T1-weighted dataset at 0.7 mm isotropic voxel resolution. The T1-weighted image was acquired using a magnetization-prepared rapid gradient echo (MP2RAGE) sequence (320 slices; TR = 5000 ms, TE = 2.47 ms; TI_1_/TI_2_ = 800/2750 ms; matrix size = 240×320×320; GRAPPA factor = 3) [41]. Additionally, a T2*-weighted anatomical dataset was acquired during the first session but was not used in the present study.

Functional MRI (fMRI) data were acquired with a 2D gradient-echo (GE) simultaneous multi-slice (SMS), multiband (MB) echo planar imaging (EPI) sequence optimised for blood-oxygen-level-dependent (BOLD) contrast [42, 43]. The acquisition parameters were as follows: TR = 1800 ms, TE = 26 ms, flip angle = 67°, 46 axial slices, 0.8 mm isotropic voxels, multi-band acceleration factor (MB) = 2, and GRAPPA factor = 3. Slice placement was adjusted to ensure adequate coverage of the auditory cortex. During scanning, participants passively listened to auditory pure-tone stimuli as part of the auditory localizer paradigm.

Session 2: fMRI data were collected using the same acquisition parameters as in Session 1. To ensure consistent anatomical coverage across sessions, slice positioning was automatically aligned to the first session using a vendor-provided alignment procedure based on a previously saved anatomical reference. Participants passively listened to stochastic auditory sequences during this session.

### 4.3 Calibration of Auditory Loudness Waveform

Prior to the experiment, participants completed a loudness calibration procedure to perceptually equalise the loudness of all stimulus frequencies. While positioned in the scanner with in-ear earphones in place (Sensimetric Inc., s15 model), participants were presented with pure-tone test sounds and adjusted the intensity of each tone to match the perceived loudness across frequencies.

Nine calibration tones, logarithmically spaced between 200 Hz and 6,000 Hz, were used to construct an individual equal-loudness contour for each participant [44]. Intermediate values were interpolated using piecewise cubic interpolation to generate a continuous equal-loudness curve. This curve was then used to adjust the intensity of all tones throughout the experiment, ensuring uniform perceived loudness across the entire stimulus frequency range.

### 4.4 Population Receptive Field Mapping of the Auditory Cortex

In the first session, population receptive field (pRF) mapping was performed to estimate voxel-wise frequency preferences and tuning widths across the auditory cortex. These frequency-selective profiles were used to parameterise voxel-specific models of the sound sequences presented in the second session. This allowed us to examine repetition suppression and expectation effects in a content-dependent manner.

#### 4.4.1 Quasi-Random Tone Sequences

We measured fMRI responses to sequences of pure tones presented in a quasi-random order to estimate population receptive fields (pRFs; [22, 21]). Auditory stimuli were generated in MATLAB (The MathWorks, Natick, MA) with the Psychophysics Toolbox extension [45, 46, 47] and delivered via MRI-compatible in-ear headphones (Sensimetric Inc., s15 model) at a 48,000 Hz sampling rate.

Each of the 240 tones was presented once per run, with a fixed inter-tone interval (ITI) of 400 milliseconds. The presentation order was quasi-randomised separately for each run, with the constraint that successive tones differed by at least one octave in frequency. This constraint reduced temporal correlations between neighbouring frequencies, limiting neural adaptation and improving the fidelity of pRF estimates.

To promote hemodynamic recovery and enable accurate baseline modeling, silent periods were interspersed throughout the sequence. Following every block of 80 tone presentations, a 14.4-second silent interval was inserted to allow estimation of baseline responses in the presence of scanner noise alone [48]. In addition, brief silent gaps were inserted on 5% of trials within the tone sequence, introducing these short interruptions further promoted recovery and mitigate stimulus-driven adaptation effects. Each run began and ended with 12.6 seconds of silence to stabilise baseline estimation.

Session one included six pRF mapping runs, each lasting 8 minutes and 20 seconds. For participant nine, one run was excluded due to data corruption, leaving five usable runs. This reduction did not substantially impact the quality of the resulting pRF estimates.

#### 4.4.2 Population Receptive Field Estimation

We estimated frequency-selective population receptive fields (pRFs) using a two-stage procedure. First, single beta weights were computed for each voxel, reflecting its response to each of the 240 acoustic frequencies across all six experimental runs. Second, we applied a permutation-based model grid search to derive voxel-specific pRF estimates and mitigate selectivity biases in low-SNR regions [49].

fMRI responses were modeled using a canonical double-gamma hemodynamic response function (HRF) with standard parameters (time to peak: 4 s, time to undershoot: 16 s, positive-to-negative ratio: 6, dispersion: 1). This HRF was convolved with the stimulus time series, and beta weights were estimated voxelwise across the full frequency range (200 Hz to 6 kHz). The resulting set of 240 beta weights per voxel served as the input for subsequent pRF estimation.

Candidate pRFs were defined as Gaussian response profiles on a log_2_-transformed frequency axis. Centre frequencies (*µ*) were sampled in equal steps across 240 log_2_-bins, and tuning widths were parameterised using full-width at half-maximum (FWHM) values ranging from 0.5 to 2.0 octaves. For each candidate model, a Gaussian response profile was constructed:

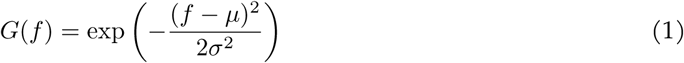

where *f* is the frequency in log_2_ space, *µ* denotes the center frequency, and *σ* is the standard deviation derived from the specified FWHM.

Following an initial grid search over all candidate pRFs, we implemented a permutation-based refinement to correct for biases in model selection, particularly the tendency to overestimate sharp tuning in low-SNR voxels. The permutation-based model grid search, as described in [49], uses a null distribution to estimate the likelihood of selecting each grid point by chance. For each voxel, 1,000 permutations were performed by randomising the frequency labels of the beta weights. At each permutation, we computed the correlation between the observed voxel responses and the predicted responses from each grid point. The optimal model was defined as the one with the lowest probability of occurrence under the null hypothesis, calculated as − log(*p*)across all permutations. This procedure improved the reliability of pRF estimates in noisy regions without distorting the estimates in voxels with high signal quality. The resulting pRF maps provided robust voxel-wise estimates of preferred frequency and tuning width across the auditory cortex.

### 4.5 Presentation of Stochastic Sequences

In session 2, auditory sequences consisting of pure-tones sampled probabilistically from mixtures of two Gaussian distributions, each defined on a log_2_-frequency axis. The sequences were drawn from one of four distinct Gaussian pairs, with center frequencies ranging from 500 Hz to 3,000 Hz: (500/1392 Hz), (646/1790 Hz), (834/2323 Hz), and (1,078/3,000 Hz). Within each pair, the Gaussian centers were separated by 1.48 octaves, ensuring that tones from the two distributions were perceptually distinguishable. Each Gaussian had a fixed bandwidth of 1/3 octave (FWHM), and tones were sampled at a resolution of 1/12th octave, yielding uniformly spaced candidate frequencies within each distribution.

For participants one and two, experimental session two consisted of 12 runs; for all other participants, the session included 10 runs to reduce overall scanning time and keep session duration within a two-hour limit. Each run was subdivided into three blocks, and each block comprised 10 stable periods. Within each stable period, tones were sampled with fixed probabilities from the two underlying Gaussian distributions, preserving local statistical regularity. The relative probability of sampling from Distribution A versus Distribution B was manipulated across stable periods, following a logistic transformation to yield values ranging from 5% to 95%. Fifteen distinct probability steps were used, spanning strongly biased conditions (e.g., 95% A / 5% B) to near-balanced mixtures (e.g., 55% A / 45% B). These probabilities were assigned pseudo-randomly across stable periods, ensuring broad coverage of the probability space while avoiding immediate repetition of extreme values. As a result, transitions between stable periods introduced shifts in the underlying statistical structure of the tone sequences. Some of these transitions were perceptually salient (e.g., a change from 95% sampling from Distribution A to 95% sampling from Distribution B), while others were subtler (e.g., 95% sampling from Distribution A to 90% sampling from Distribution A). Because sampling was probabilistic, the perceptual clarity of these transitions varied both within and across runs.

Each tone within the stochastic sequences was presented for 200 ms, followed by a 50 ms inter-tone interval, yielding a total stimulus onset asynchrony (SOA) of 250 ms. Within each stable period, 48 tones were presented over 12 seconds, after which the sampling probability was updated for the subsequent stable period. To facilitate hemodynamic recovery and support baseline estimation, each block concluded with a 14.4-second silent interval. In addition, each run began and ended with a 9-second silent period to allow the hemodynamic response to stabilize. The total duration of each run was approximately 7.5 minutes.

### 4.6 Functional Data Analysis

#### 4.6.1 Preprocessing and Alignment of Functional Data

Functional data preprocessing included slice timing correction, motion correction, temporal filtering, and distortion correction. Unless otherwise specified, all steps were performed using BrainVoyager version 22.2 (Brain Innovation, The Netherlands; [50]). Slice timing correction was performed using sinc interpolation to account for differences in slice acquisition times, followed by 3D motion correction using rigid-body alignment to estimate head movement across six degrees of freedom (three translations and three rotations). Within each run, all volumes were aligned to the first volume of that run. Subsequently, each run was aligned to the first run of the first session. All transformations were applied in a single interpolation step. To remove low-frequency drifts, a high-pass filter with a cutoff of 7 cycles per run was applied. Geometric distortions from phase encoding were corrected using reversed phase-encoded images and FSL’s Topup algorithm [51, 52]. Non-linear distortion correction was then performed using Advanced Normalisation Tools (ANTs) to reduce geometric inconsistencies across sessions and enhance inter-session alignment [53]. Finally, functional images were aligned to the upsampled anatomical image using boundary-based registration (BBR), ensuring precise anatomical correspondence [54].

#### 4.6.2 Region of Interest (ROI) Selection

Anatomical data processing involved defining five regions of interest (ROIs) on each individual’s cortical surface, separately for each hemisphere. Based on macroanatomical landmarks [55, 33], we delineated Heschl’s gyrus (HG), Planum polare (PP), Planum temporale (PT), the anterior superior temporal gyrus (aSTG), and the posterior superior temporal gyrus (pSTG) (see Figure **S7**). These surface-based ROIs were subsequently projected into volumetric space, extending 3 mm in both directions from the mid-gray matter surface. For further analyses, the ROIs from both hemispheres were concatenated to form a single bilateral ROI.

#### 4.6.2 Layer Selection

The cerebrospinal fluid (CSF)–grey matter and gray matter–white matter boundaries were identified in volume space using BrainVoyager’s deep learning-based segmentation algorithm (Tiramisu; [56]). These boundaries were subsequently inspected and manually refined using 3D Slicer [57].

To estimate voxel-wise depth within the grey matter, we employed an equi-volume approach, which accounts for cortical curvature by ensuring equal volume fractions across depth levels [34]. In volume space, the grey matter was partitioned into three depth compartments.

Because depth estimation was conducted in anatomical space (0.4 mm isotropic) while functional data were analysed at their native resolution (0.8 mm isotropic), projecting the data into functional space caused some voxels to span multiple depth compartments. To address this, each voxel was assigned to a single depth bin based on its majority volume overlap across the three compartments. Specifically, we calculated the proportion of each voxel’s overlap with the deep, middle, and superficial layers after projection, and assigned it exclusively to the compartment with the highest overlap.

### 4.7 Computational Modelling

#### 4.7.1 Frequency-Specific Stimulus Drive Model

To estimate the frequency-specific stimulus drive of each voxel, we employ a tuning-based activation model derived from the population receptive field (pRF) measurements. Specifically, for each tone of frequency *x*_*t*_ presented, we computed the predicted response of voxel*v*as the value of its Gaussian tuning curve at that frequency:

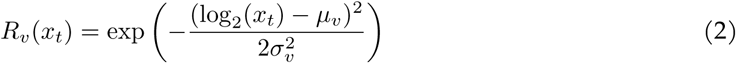

Where *log*_2_(*x*_*t*_) represents the frequency of the tone in *log*_2_-space, *µ*_*v*_ is the voxel’s peak sensitivity in the same space, and *σ*_*v*_ reflects the tuning bandwidth of voxel *v*. This Gaussian function models the expected response amplitude based on how close the tone is to the voxel’s frequency preference.

#### 4.7.2 Long-trace Repetition Suppression Model

We implemented a double-exponential model of long-trace repetition suppression, based on evidence that brief sensory exposure can induce persistent, stimulus-specific suppression in early sensory cortex. The model assumes that the influence of a stimulus presented *n* trials ago decays over time via two exponential components, reflecting distinct short- and long-lived suppression processes. Formally, the repetition suppression factor *f* (*n*) is defined as:

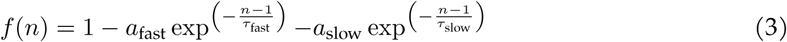

where *a*_fast_ and *a*_slow_ determine the amplitudes of the fast and slow suppression components, and *τ*_fast_ and *τ*_slow_ are their respective time constants, expressed in units of trials. We adopted parameter values from a prior neurophysiological study in primary visual cortex (V1) [8], setting *a*_fast_ = 13.99% with *τ*_fast_ = 0.85 trials, and *a*_slow_ = 3.45% with *τ*_slow_ = 6.82 trials. These values capture two temporally distinct mechanisms: a rapid suppression phase and a slower, more sustained suppression process.

To model cumulative suppression across both short and long timescales, each voxel’s effective response to a new stimulus was weighted by the combined influence of the preceding ten trials. This was computed using an *n*-back decay array derived from the double-exponential formulation. Within a tuning-dependent framework, this suppression factor was applied multiplicatively to each voxel’s frequency-tuned Gaussian activation curve, modulating the amplitude of its response based on stimulus history.

This approach generates a response grid spanning frequency and tuning width dimensions, providing repetition suppression estimates over time for each grid position and ensuring that suppression is modelled across voxels with different tuning preferences.

#### 4.7.3 D-REX Model

The Dynamic Regularity Extraction (D-REX) model employs Bayesian sequential prediction with perceptual constraints, primarily focusing on memory (*m*), to simulate the brain’s processing of sound sequences over time [31]. Memory (*m*) restricts the number of prior time points considered; here, we set *m* to remain within the boundaries of a block. The model continuously predicts the distribution of the next time point (*x*_*t*+1_) based on previous observations (*x*_1:*t*_) by estimating local statistics (*θ*), corresponding to the sample mean and variance of the input.

In addition to these features, the model incorporates a Gaussian Mixture Model (GMM) to represent multiple components within the sound sequences. For our stimuli specifically, we constrained the maximum number of components to two, corresponding to the two centre frequencies present. To manage the formation of new components within the GMM, a threshold parameter, *β*, was set to 0.2. This *β* value, scaled by predictive probabilities, further constrains the overall stimulus space.

The model accounts for possible changes in parameters (*θ*) by generating hypotheses across varying run lengths (*r*_*t*_) and integrating these to predict future observations:

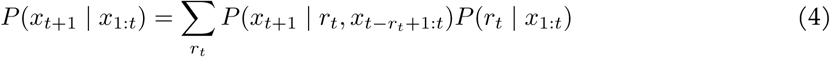

The primary output of the model is surprisal (*S*_*t* +1_), representing the discrepancy between the predicted probability and the actual observation:

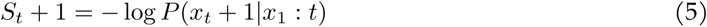

Lower probability events yield higher surprisal, while zero surprisal corresponds to perfectly predicted events. Additionally, the model computes precision, defined as the inverse of the variance across the full Gaussian mixture predictive distribution, thereby quantifying the confidence in predictions over time:

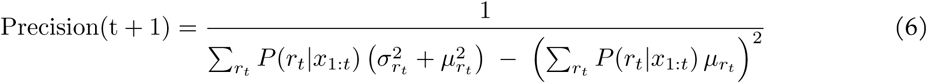

Here, *P* (*r*_*t*_ | *x* _1: *t*_) is the posterior probability of a run length *r*_*t*_ given observations up to time *t*, while 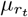 and 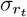 denote the predicted mean and standard deviation, respectively, of the next stimulus under run length *r*_*t*_.

Beyond computing surprisal and precision for each observed stimulus, the model estimates the prior probability of each frequency across the stimulus space within the inferred predictive landscape. This continuous probability distribution explicitly reflects the model’s expectations based on previously encountered stimuli, providing a structured representation of auditory regularities utilised for tuning-dependent analyses.

#### 4.7.4 Prediction Error Estimation

To quantify the mismatch between model-based expectations and stimulus-driven activation, we define a voxel-wise prediction error metric. At each time point *t*, prediction error for voxel *v* is computed as the absolute difference between the expected activation based on the model’s prior and the stimulus drive response to the observed stimulus:

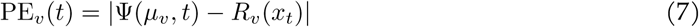

Here, *R*_*v*_ (*x*_*t*_) is the stimulus-driven activation of voxel *v* based on its tuning function (see2), and Ψ (*f, t*) denotes the prior probability assigned by the D-REX model to frequenc y fat time *t*. Thus, Ψ (*µ*_*v*_, *t*) reflects the model’s expected probability at the voxel’s preferred frequency. This measure captures the magnitude of prediction error as the discrepancy between expected and observed activation in frequency space.

### 4.8 Stimulus Modeling

Stimulus modeling was implemented on a tone-by-tone basis to construct regressors corresponding to each tone presented during the experiment. These regressors were temporally aligned with presentation logs, ensuring accurate representation of tone onsets and offsets in the time domain. All tones had a fixed duration of 200 ms and were separated by a 50 ms inter-stimulus interval (ISI), establishing a modelling resolution of 50 ms. Each regressor was convolved with a canonical double-gamma hemodynamic response function (HRF), peaking at 4 s with an undershoot at 16 s — consistent with the HRF employed during population receptive field (pRF) mapping in session one. Because the functional MRI (fMRI) repetition time (TR) far exceeded the resolution of the stimulus model, convolved regressors were resampled to the TR timescale, ensuring alignment with fMRI acquisition while preserving their temporal structure.

We specified three primary models, each featuring a distinct combination of regressors capturing different neural response aspects. (1) The stimulus drive model included voxel-specific raw stimulus activation based on estimated tuning curves and a simple on/off regressor marking tone occurrences. (2) The repetition suppression model integrated regressors reflecting adaptation-sensitive responses derived from the repetition suppression model described previously. (3) The expectation model incorporated regressors from probabilistic modelling, including prior probabilities, surprisal, precision, and prediction error, as defined previously.

Voxelwise regression analyses followed a tuning-dependent framework, explicitly aligning regressors with voxel-specific tuning properties. Regressors for stimulus drive, repetition suppression, prior probabilities, and prediction error were linked directly to each voxel’s frequency preference, while surprisal and precision were applied uniformly across voxels.

To enhance run-wise comparability and reduce high-frequency noise, fMRI responses were z-scored and temporally smoothed per run using a Gaussian filter (sigma = 1), consistent with the temporal profile of the HRF.

### 4.9 Cross-validated Regression Approach

We used leave-one-out (LOO) cross-validation to evaluate model generalisation. In this approach, data from one run was withheld while the model was trained on the remaining runs, and this process was repeated for all runs.

In addition to cross-validation, we performed an additional regression pass on the full dataset without data splits. This step served two purposes: (1) selecting voxels based on the full model fit and (2) generating functional maps (f-maps) to visualise overall performance. To aid in voxel selection, we computed uncorrected significance levels (p-values) from these f-maps.

To summarise model performance, we reported the median prediction accuracy on the held-out run across folds. Using the median rather than the mean provided a more robust measure, reducing the impact of single fold outliers.

### 4.10 Variance Partitioning

To quantify the *unique* and *overlapping* cross-validated variance explained by Repetition Suppression and Expectations relative to the Stimulus Drive baseline, we relied on *variance partitioning* using Set Theory.

We employed a similar approach to that described by [32]. *Set Theory* provides a mathematical framework for defining and analysing unions and intersubsections between sets. For three sets (*A, B, C*), we can express the unique contributions and their intersubsections as:

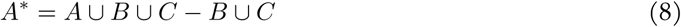

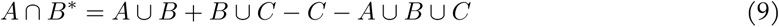

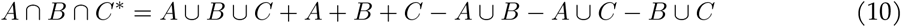

Using the additive nature of sets, we calculated the unique contributions of each set (*A*^*^, *B*^*^, *C*^*^) and the unique inter-subsections between sets (*A* ∩ *B*^*^, *A* ∩ *C*^*^, *B* ∩ *C*^*^, *A* ∩ *B* ∩ *C*^*^) based on the formulas in Equations8,9, and10.

For our analyses, we capitalised on the linear additive property of variance explained ( *R*^2^), allowing us to calculate the *unique* and *overlapping* variance contributions of Repetition Suppression and Expectations relative to the Stimulus Drive Model.

Formally, we calculated the cross-validated variance explained for the following models: *(Stimulus Drive), (Repetition Suppression), (Expectations), (Stimulus Drive* ∪ *Repetition Suppression), (Stimulus Drive* ∪ *Expectations), (Repetition Suppression* ∪ *Expectations)*, and *(Stimulus Drive* ∪ *Repetition Suppression* ∪ *Expectations)*. Union models included regressors from all models within the specified union.

### 4.11 Statistical Testing

Statistical analyses were conducted across participants. Given the relatively small sample size (N=10), we employed non-parametric bootstrap t-tests. This approach involved resampling a null distribution with a mean of zero, achieved by centring the data through mean removal. We then estimated the likelihood of obtaining a t-value at least as extreme as the observed t-value across 10,000 bootstrap iterations.

P-values were computed without assuming symmetry, using an equal-tailed approach as recommended by Rousselet et al. (2019). Confidence intervals, both in figures and text, were also derived via bootstrapping. The primary statistical tests assessed the cross-validated unique variance explained by each explanatory model, over and above the variance explained by alternative models.

Statistical tests were conducted separately for each region of interest (ROI). In addition, we conducted pairwise comparisons between cortical layers (deep, middle, superficial) using paired-sample, non-parametric bootstrap t-tests. These layer-wise comparisons followed the same resampling procedure as described above.

To control for multiple comparisons, false discovery rate (FDR) correction was applied using the Benjamini–Hochberg procedure [58]. For the primary tests assessing unique variance explained by each model, FDR correction was applied across five comparisons (corresponding to the five ROIs). For the pairwise layer-wise comparisons (deep vs middle, deep vs superficial, middle vs superficial), FDR correction was applied across fifteen comparisons in total (3 comparisons per ROI×5 ROIs).

## 4.12 Acknowledgements

We thank Lonike Faes and Mahdi Enan for their insightful scientific discussions. This work was supported by the European Research Council (ERC) under the European Union’s Horizon 2020 research and innovation programme (grant agreement No. 101001270).

## 4.13 Author Contributions

Conceptualisation: JJGvH. Data wrangling and preprocessing: JJGvH. Formal analysis: JJGvH. Statistical analysis and visualisation: JJGvH. Supervision: FdM, SAK, FPdL. Initial draft: JJGvH. Final draft: JJGvH, SAK, FPdL, FdM.

## 4.14 Competing Interests

The authors declare no competing interests.

## 4.15 Data Availability

Anatomical and functional MRI data are available on OpenNeuro (openneuro.org/datasets/ds006928). Questions about the dataset can be directed to the corresponding author, Jorie van Haren.

## 4.16 Code Availability

All analysis scripts and modeling code are available on GitHub (github.com/mesoScopic-Computational-AuditioN-lab). Additional details can be requested from the corresponding author, Jorie van Haren.

## Notes

### Competing Interest Statement

The authors have declared no competing interest.

### Summary of Updates

This version of the manuscript includes minor corrections to grammatical errors and typographical mistakes. The Data and Code Availability sections have been updated to include final repository links and descriptions.

