## Supplementary materials for "Distinct Roles of Deep and Superficial Cortical Layers in Tone Prediction, Comparison, and Adaptation in Human Auditory Cortex"

### A Supplement

#### A.1 Widespread Stimulus Drive and Repetition Suppression Throughout Auditory Cortex, with Regionally Selective Expectation Effects

To assess how stimulus drive, repetition suppression, and expectation-related signals are distributed across the auditory hierarchy, we quantified their unique explained variance within anatomically defined regions of interest (ROIs) spanning primary auditory cortex (Heschl's gyrus, HG), belt regions (Planum Polare, PP; Planum Temporale, PT), and associative auditory cortex along superior temporal gyrus (anterior STG, aSTG; posterior STG, pSTG).

In primary auditory cortex (Heschl's gyrus, HG), both stimulus drive (5.3% of the cumulative explained variance, hereafter abbreviated as EV; FDR-corrected across five ROIs, $p_{\mathrm{FDR}} = 2.4\mathrm{e}{-2}$) and repetition suppression (8.9% EV; $p_{\mathrm{FDR}} = 1.7\mathrm{e}{-4}$) explained significant and unique portions of the observed fMRI time courses. In contrast, expectation (3.3% EV; $p_{\mathrm{FDR}} = 0.11$) did not contribute significantly beyond what was already captured by stimulus drive and adaptation (see figure S1a). This suggests that in HG, expectation components are not readily distinguishable from more parsimonious explanations grounded in frequency selectivity and recent acoustic history.

Moving outward to auditory belt regions, differences in the relative contributions of stimulus drive, repetition suppression, and expectation became more apparent. Within Planum Polare (PP), each of the three factors explained a significant amount of unique variance: expectation emerging as the strongest contributor (23.2% EV; $p_{\mathrm{FDR}} < 1\mathrm{e}{-5}$), followed by stimulus drive (20.1% EV; $p_{\mathrm{FDR}} < 1\mathrm{e}{-5}$) and repetition suppression (17.1% EV; $p_{\mathrm{FDR}} = 8.3\mathrm{e}{-5}$) (see figure S1b). Compared to HG, all three components contributed more distinctly to PP, indicating a clearer functional separation between frequency selectivity, recent stimulus history, and predictive information, while overall model fit remained comparable across the two regions.

In contrast to PP, the profile in Planum Temporale (PT) aligned more closely with that of HG, with significant unique contributions from stimulus drive (6.5% EV; $p_{\mathrm{FDR}} = 1.2\mathrm{e}{-2}$) and repetition suppression (9.0% EV; $p_{\mathrm{bonf}} = 3.4\mathrm{e}{-4}$), but no significant unique variance explained by expectation (4.8% EV; $p_{\mathrm{FDR}} = 0.11$) (see figure S1c). As in HG, variance linked to expectation in PT may be explained by stimulus drive and repetition-based processes, reflecting a less differentiated encoding of predictive information. Notably, across both HG and PT, repetition suppression consistently accounted for a distinct portion of the variance, pointing to its distributed involvement throughout early and intermediate auditory regions.

In associative auditory regions along the superior temporal gyrus (STG), expectation effects became more prominent, alongside persistent significant contributions of stimulus drive and repetition suppression. Both anterior-STG (aSTG) and posterior-STG (pSTG) showed significant unique contributions of each of the modeled processes. In aSTG, stimulus drive (14.3% EV; $p_{\mathrm{FDR}} < 1\mathrm{e}{-5}$), repetition suppression (13.6% EV; $p_{\mathrm{FDR}} < 1\mathrm{e}{-5}$), and expectation (16.2% EV; $p_{\mathrm{FDR}} = 8.3\mathrm{e}{-3}$) each explained meaningful unique variance (see figure S1d). Similarly, in pSTG, stimulus drive (13.2% EV; $p_{\mathrm{FDR}} = 3.5\mathrm{e}{-3}$), repetition suppression (14.9% EV; $p_{\mathrm{FDR}} = 8.3\mathrm{e}{-5}$), and expectation (11.3% EV; $p_{\mathrm{FDR}} = 4.0\mathrm{e}{-4}$) contributed significantly, reflecting a more balanced distribution across the three processes (see figure S1e). This emergence of distinct contributions suggests that stimulus drive, repetition suppression, and expectation capture different functional aspects of the response along the STG, with reduced overlap. Notably, the inclusion of the expectation component explained aspects of the response that were not accounted for by stimulus drive or recent acoustic history, indicating that at this level, predictive information must be integrated to better capture the structure of the response.

#### A.2 Consistent Deep-Layer Bias of Prediction Errors Across the Auditory Hierarchy and Region-Specific Bias of Priors

In this supplementary analysis, we reformulated the model to subsume repetition suppression into the stimulus drive baseline, thereby allowing priors and prediction errors to be partitioned as distinct sources of variance. This adjustment enabled us to track their unique contributions across cortical

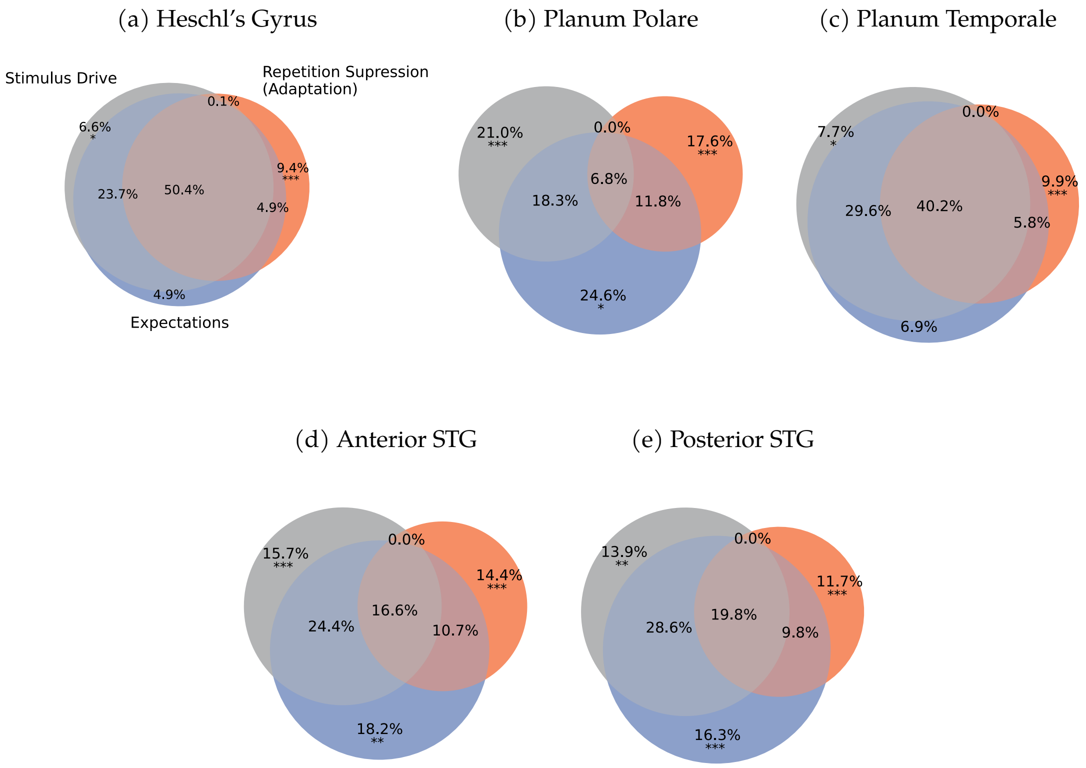


**Figure S1**: **Stimulus drive and repetition suppression effects are widespread across auditory cortex, while expectation-related variance emerges selectively in belt- and associative regions.** Venn diagrams summarize voxelwise BOLD variance explained by stimulus drive, repetition suppression, and expectations. For each ROI, set-theoretic variance partitioning shows the proportional breakdown of total explained variance (EV) into unique and shared components. For an overview of the ROI selection, see Figure S7. Stars indicate significance levels of the cross-validated unique variance explained (one-sample bootstrap t-tests, FDR-corrected across ROIs): $p_{\mathrm{FDR}} < 0.05$ (*), $p_{\mathrm{FDR}} < 0.01$ (**), $p_{\mathrm{FDR}} < 0.001$ (***). a HG: strong unique contributions from stimulus drive and repetition suppression; expectation did not explain additional variance. b PP: clear separation between stimulus drive, adaptation, and expectation, each explaining distinct portions of EV. c PT: like HG, with robust stimulus drive and repetition suppression; expectation again non-significant. d aSTG and e pSTG: all three components contributed uniquely, with expectation explaining additional variance beyond stimulus drive and adaptation.

depth. However, this reformulation necessarily precluded estimating their shared variance with stimulus drive or with each other, and thus isolates the components at the cost of losing information about their joint contributions.

Prediction errors uniquely contributed to deep-layer responses across the auditory cortex, with significant laminar biases in all five ROIs (see figure S2). In Heschl's gyrus (HG), error explained more unique variance in deep compared to both middle (FDR-corrected across 15 comparisons spanning 5 ROIs and 3-layer contrasts, $p_{\mathrm{FDR}} < 1.0\mathrm{e}{-5}$) and superficial layers ($p_{\mathrm{FDR}} = 2.3\mathrm{e}{-2}$). In Planum Polare (PP), a similar pattern emerged, with more variance explained in deep compared to superficial layers ($p_{\mathrm{FDR}} = 1.3\mathrm{e}{-2}$). In Planum Temporale (PT), error-related variance exhibited a consistent gradient across depth, with greater unique variance explained between deep and superficial layers ($p_{\mathrm{FDR}} = 2.3\mathrm{e} - 2$), deep and middle layers ($p_{\mathrm{FDR}} = 7.0\mathrm{e} - 3$), and middle and superficial layers ($p_{\mathrm{FDR}} = 2.3\mathrm{e} - 2$). In anterior superior temporal gyrus (aSTG), error signals were again stronger in deep versus superficial layers ($p_{\mathrm{FDR}} = 1.5\mathrm{e}{-3}$) and in middle versus superficial layers ($p_{\mathrm{FDR}} = 6.8\mathrm{e}{-3}$). Finally, in posterior STG (pSTG), more unique variance was explained in deep compared to superficial layers ($p_{\mathrm{FDR}} = 1.8\mathrm{e}{-3}$) and in deep compared to middle layers ($p_{\mathrm{FDR}} = 1.5\mathrm{e}{-2}$).

Prior-related activity exhibited weaker laminar-specific effects, with significant differences between layers limited to higher-order regions (see figure S2). In Planum Temporale (PT), priors accounted for more unique variance in deep compared to superficial layers ($p_{\mathrm{FDR}} = 5.7\mathrm{e}{-3}$). Similarly, in anterior STG (aSTG), priors showed a deep-layer preference, with greater unique variance explained in deep compared to superficial layers ($p_{\mathrm{FDR}} = 3.6\mathrm{e}{-2}$). No other regions showed reliable depth effects. These focal findings suggest that priors engage infragranular layers selectively in higher-order auditory areas, while remaining more diffusely expressed elsewhere.

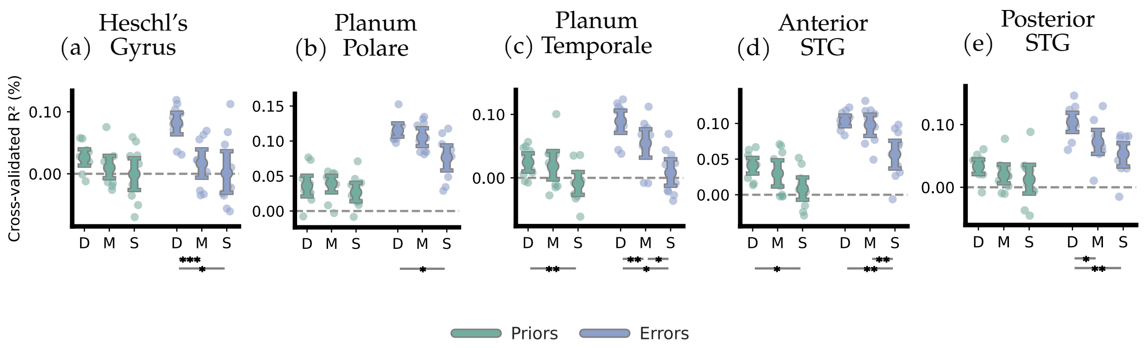


**Figure S2**: **Laminar profiles of priors and prediction errors: consistent deep-layer bias for errors, focal deep effects for priors.** Each panel shows the cross-validated unique variance (variance explained by each model component beyond what could be attributed to stimulus drive and repetition suppression) accounted for by prior probability and prediction error across cortical layers. D, M, and S denote deep, middle, and superficial layers, respectively. Results are shown for five auditory regions of interest (ROIs): a Heschl's Gyrus (HG), b Planum Polare (PP), c Planum Temporale (PT), d anterior Superior Temporal Gyrus (aSTG), and e posterior STG (pSTG). Prediction error-related activity consistently exhibited a deep-layer bias across all ROIs, while prior-related activity showed a more restricted laminar profile, with significant depth effects in PT and aSTG only. Stars indicate significance levels of the cross-validated unique variance explained across depth (paired-sample bootstrap t-tests, FDR-corrected across all layer pair comparisons and ROIs): $p_{\mathrm{FDR}} < 0.05$ (*), $p_{\mathrm{FDR}} < 0.01$ (**), $p_{\mathrm{FDR}} < 0.001$ (***). Error bars indicate bootstrapped 95% confidence intervals.

#### A.3 Progressive Shortening of Repetition Suppression Window from Heschl's Gyrus to Higher Order Regions

To characterize how repetition suppression dynamics evolve across the auditory hierarchy, we modeled suppression at two timescales using fast ($a_{\mathrm{fast}}, \tau_{\mathrm{fast}}$) and slow ($a_{\mathrm{slow}}, \tau_{\mathrm{slow}}$) exponential decay components, and systematically sampled a grid of suppression profiles to estimate how much and how long prior stimulation influenced current responses in each cortical region. The fast component was varied across six positions, with $a_{\mathrm{fast}}$ decreasing from 20% to 4% and $\tau_{\mathrm{fast}}$ ranging from 1 to 0.01 trials. The slow component was varied across four positions, with $a_{\mathrm{slow}}$ increasing from 3% to 12% and $\tau_{\mathrm{slow}}$ ranging from 7 to 3 trials. For each grid point, we generated an repetition suppression regressor and quantified its fit to the voxel timecourse using $R^2$. The best-fitting grid position thus provided voxel-wise and ROI-based estimates of $a_{\mathrm{fast}}, \tau_{\mathrm{fast}}, a_{\mathrm{slow}}, \tau_{\mathrm{slow}}$. These estimates were obtained from the full dataset (no cross-validation) due to computational cost. Mapping these best-fit parameters onto the cortical surface revealed distinct spatial patterns of fast and slow suppression, offering insight into how stimulus history shapes neural responses across the auditory hierarchy (see Fig. S3).

In Heschl's gyrus (HG), the best-fitting grid point contained the strongest and most persistent fast component ($a_{\mathrm{fast}} = 0.10, ; \tau_{\mathrm{fast}} = 0.41$ trials; 95% CI, bootstrapped across participants: $a_{\mathrm{fast}} = [0.067; 0.108]$, $\tau_{\mathrm{fast}} = [0.189; 0.427]$). This parameterization generates an immediate 10% drop in response magnitude that decays only gradually, with about 37% of the initial suppression still present at the next tone. Combined with a small but long-lasting slow component ($a_{\text{slow}} = 0.03; \tau_{\text{slow}} = 7$ trials; 95% CI: $a_{\text{slow}} = [0.030; 0.057]$, $\tau_{\text{slow}} = [5.8; 7.0]$), this pattern suggests that stimulus history exerts the strongest and most prolonged influence at this early cortical stage.

In Planum Polare (PP), the repetition-suppression window shortened considerably: the fast magnitude decreased to $a_{\text{fast}} = 0.04$ and the time constant collapsed to $\tau_{\text{fast}} = 0.01$ trials (95% CI: $a_{\text{fast}} = [0.046; 0.073]$, $\tau_{\text{fast}} = [0.030; 0.230]$), so virtually all fast suppression had dissipated by the time the next stimulus occurred. However, the slow component remained unchanged ($a_{\text{slow}} = 0.03, ; \tau_{\text{slow}} = 7$ trials; 95% CI: $a_{\text{slow}} = [0.030; 0.030]$, $\tau_{\text{slow}} = [7.0; 7.0]$), indicating that although sensitivity to immediate repetitions was reduced, slower integrative effects continued to shape responses over longer timescales.

In Planum Temporale (PT) and both the anterior and posterior superior temporal gyrus (aSTG, pSTG), intermediate fast parameters emerged ($a_{\text{fast}} = 0.07, ; \tau_{\text{fast}} = 0.21$ trials; 95% CI: PT: $a_{\text{fast}} = [0.052; 0.082]$, $\tau_{\text{fast}} = [0.090; 0.290]$; aSTG/pSTG: $a_{\text{fast}} = [0.046; 0.070]$, $\tau_{\text{fast}} = [0.050; 0.210]$). These values recover roughly twice as quickly as in HG but remain considerably slower than in PP. Across all regions, the slow component was remarkably stable ($a_{\text{slow}} \approx 0.03; \tau_{\text{slow}} \approx 7$ trials; 95% CI: PT: $a_{\text{slow}} = [0.030; 0.066]$, $\tau_{\text{slow}} = [5.399; 7.000]$; aSTG/pSTG: $a_{\text{slow}} = [0.030; 0.030]$, $\tau_{\text{slow}} = [7.0; 7.0]$), suggesting that long-range stimulus history is represented uniformly throughout the hierarchy. What differentiates cortical stages is primarily the speed and strength of the fast component: as one moves away from HG, the immediate suppression becomes progressively smaller and its recovery accelerates, effectively contracting the repetition-suppression integration window for fast adaptation, while the slow component maintains a consistent background influence across regions.

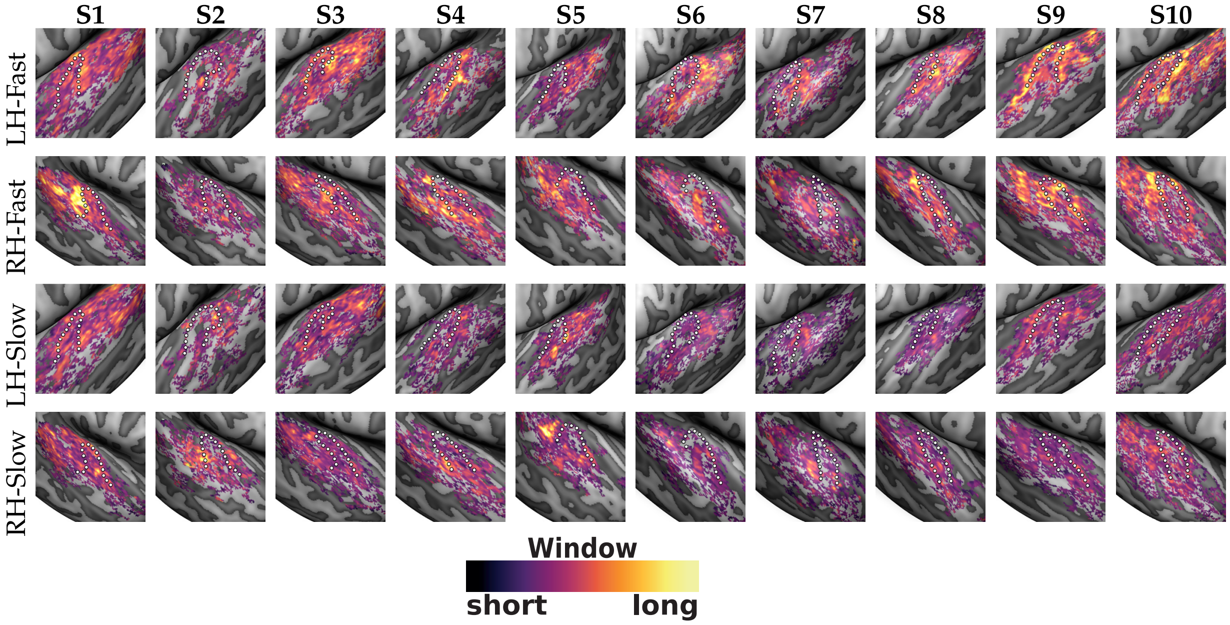


**Figure S3**: **Individual subject maps of fast and slow repetition suppression components.** Rows show voxelwise maps of the estimated fast and slow components of the repetition suppression window, separately for left (LH) and right (RH) hemispheres. Columns correspond to individual participants (S1–S10). Maps were manually masked to auditory cortex and thresholded at $p < .05$ (uncorrected) fit of the repetition suppression model. The dotted white line denotes the anatomical border of Heschl's gyrus. Suppression windows were estimated by fitting a grid of parameter combinations: the fast component varied from $a_{\text{fast}} = 20\%$ to 4% and $\tau_{\text{fast}}$ from 1 to 0.01 trials; the slow component from $a_{\text{slow}} = 3\%$ to 12% and $\tau_{\text{slow}}$ from 7 to 3 trials. For each voxel, the color reflects the best-fitting parameter combination. Cooler colors indicate briefer and weaker suppression windows, while hotter colors reflect longer and more dampening profiles.

| | Deep vs Middle | | Middle vs Superficial | | Deep vs Superficial | | ROI Level | |
|---|---|---|---|---|---|---|---|---|
| | $R^2$ | $p_{\text{FDR}}$ | $R^2$ | $p_{\text{FDR}}$ | $R^2$ | $p_{\text{FDR}}$ | $R^2$ | $p_{\text{FDR}}$ |
| **Heschl's gyrus** | | | | | | | | |
| $StimulusDrive$ | 2.7e-4 | 2.1e-1 | 1.9e-4 | 3.2e-1 | 4.5e-4 | 1.3e-1 | 3.6e-4 | 2.4e-2 |
| $RepetitionSuppression$ | -7.4e-5 | 9.6e-1 | -3.2e-5 | 9.6e-1 | -1.1e-4 | 9.6e-1 | 6.1e-4 | 1.7e-4 |
| $Expectation$ | 7.9e-4 | 6.0e-4 | 1.3e-4 | 5.6e-1 | 9.2e-4 | 7.2e-3 | 2.3e-4 | 1.1e-1 |
| $StimulusDrive \cap RepetitionSuppression$ | 4.8e-5 | 8.7e-1 | -1.8e-4 | 7.0e-1 | -1.3e-4 | 7.8e-1 | 2.7e-5 | ~1.0 |
| $StimulusDrive \cap Expectation$ | -5.3e-4 | 6.8e-2 | -3.2e-4 | 1.8e-1 | -8.5e-4 | 4.6e-3 | 1.6e-3 | 5.6e-4 |
| $RepetitionSuppression \cap Expectation$ | 1.6e-4 | 3.7e-1 | 4.3e-5 | 7.1e-1 | 2.0e-4 | 1.7e-1 | 2.8e-4 | 2.4e-3 |
| $StimulusDrive \cap RepetitionSuppression \cap Expectation$ | -2.2e-3 | <1.0e-5 | -1.9e-3 | 4.9e-3 | -4.1e-4 | 1.2e-3 | 3.7e-3 | 3.5e-4 |
| **Planum Polare** | | | | | | | | |
| $StimulusDrive$ | 9.1e-5 | 3.5e-1 | 7.0e-5 | 4.7e-1 | 1.6e-4 | 3.2e-1 | 8.9e-4 | <1.0e-5 |
| $RepetitionSuppression$ | -8.1e-5 | 9.6e-1 | 1.3e-4 | 9.6e-1 | 5.2e-5 | 9.6e-1 | 7.6e-4 | 8.3e-5 |
| $Expectation$ | 9.1e-5 | 5.6e-1 | 3.5e-4 | 7.2e-2 | 4.4e-4 | 3.7e-2 | 1.0e-3 | <1.0e-5 |
| $StimulusDrive \cap RepetitionSuppression$ | -9.7e-5 | 7.0e-1 | -7.1e-5 | 7.8e-1 | -1.7e-4 | 7.0e-1 | -3.7e-4 | ~1.0 |
| $StimulusDrive \cap Expectation$ | -1.5e-4 | 1.6e-1 | -2.0e-4 | 1.2e-1 | -3.6e-4 | 6.8e-2 | 8.3e-4 | 1.7e-5 |
| $RepetitionSuppression \cap Expectation$ | 1.1e-4 | 2.3e-1 | -4.1e-5 | 7.0e-1 | 7.3e-5 | 4.1e-1 | 5.0e-4 | <1.0e-5 |
| $StimulusDrive \cap RepetitionSuppression \cap Expectation$ | -1.8e-4 | 6.6e-2 | -9.4e-4 | 4.2e-3 | -1.1e-3 | 2.7e-3 | 4.3e-4 | 2.6e-2 |
| **Planum Temporale** | | | | | | | | |
| $StimulusDrive$ | 1.5e-4 | 2.5e-1 | 2.8e-4 | 1.1e-1 | 4.2e-4 | 1.5e-1 | 4.1e-4 | 1.2e-2 |
| $RepetitionSuppression$ | 4.5e-5 | 9.6e-1 | -8.7e-6 | 9.6e-1 | 3.7e-5 | 9.6e-1 | 5.7e-4 | 3.4e-4 |
| $Expectation$ | 3.6e-4 | 5.3e-2 | 5.3e-4 | 7.4e-3 | 8.9e-4 | 7.3e-3 | 3.1e-4 | 1.1e-1 |
| $StimulusDrive \cap RepetitionSuppression$ | 2.0e-5 | 8.7e-1 | -1.1e-4 | 7.0e-1 | -8.8e-5 | 7.8e-1 | -1.8e-4 | ~1.0 |
| $StimulusDrive \cap Expectation$ | -3.0e-4 | 1.2e-1 | -8.8e-4 | 1.3e-3 | -1.2e-3 | 1.3e-3 | 1.9e-3 | <1.0e-5 |
| $RepetitionSuppression \cap Expectation$ | 2.0e-4 | 9.8e-2 | -3.1e-5 | 7.8e-1 | 1.7e-4 | 1.3e-1 | 3.4e-4 | 5.9e-3 |
| $StimulusDrive \cap RepetitionSuppression \cap Expectation$ | -1.5e-3 | 1.2e-3 | -1.9e-3 | 2.0e-3 | -3.4e-3 | 1.2e-3 | 2.8e-3 | 5.7e-4 |
| **anterior STG** | | | | | | | | |
| $StimulusDrive$ | 9.9e-5 | 3.2e-1 | 2.6e-4 | 1.1e-1 | 3.6e-4 | 1.1e-1 | 6.6e-4 | <1.0e-5 |
| $RepetitionSuppression$ | -1.5e-6 | 9.6e-1 | 4.9e-5 | 9.6e-1 | 4.7e-5 | 9.6e-1 | 6.3e-4 | <1.0e-5 |
| $Expectation$ | 1.6e-4 | 1.5e-1 | 5.3e-4 | 6.0e-4 | 7.0e-4 | 2.2e-3 | 7.5e-4 | 8.3e-3 |
| $StimulusDrive \cap RepetitionSuppression$ | 2.2e-5 | 8.7e-1 | -1.6e-5 | 8.7e-1 | 6.5e-6 | 9.7e-1 | -2.0e-4 | ~1.0 |
| $StimulusDrive \cap Expectation$ | -2.3e-4 | 9.9e-2 | -3.9e-4 | 6.8e-2 | -6.2e-4 | 1.5e-2 | 1.1e-3 | 2.0e-4 |
| $RepetitionSuppression \cap Expectation$ | 1.7e-4 | 9.8e-2 | 1.0e-4 | 9.8e-2 | 2.7e-4 | 2.1e-2 | 4.5e-4 | <1.0e-5 |
| $StimulusDrive \cap RepetitionSuppression \cap Expectation$ | -7.0e-4 | 4.6e-3 | -1.5e-3 | 2.3e-3 | -2.2e-3 | 2.0e-3 | 1.0e-3 | 7.1e-3 |
| **posterior STG** | | | | | | | | |
| $StimulusDrive$ | 2.3e-4 | 1.5e-3 | 1.2e-4 | 2.1e-1 | 3.5e-4 | 9.4e-2 | 6.5e-4 | 3.5e-3 |
| $RepetitionSuppression$ | 3.5e-6 | 9.6e-1 | 8.5e-5 | 9.6e-1 | 8.8e-5 | 9.6e-1 | 5.6e-4 | 8.3e-5 |
| $Expectation$ | 4.7e-4 | 6.0e-4 | 2.8e-4 | 1.1e-2 | 7.5e-4 | 6.0e-4 | 7.3e-4 | 4.0e-4 |
| $StimulusDrive \cap RepetitionSuppression$ | -6.4e-5 | 7.0e-1 | -3.9e-5 | 7.8e-1 | -1.0e-4 | 7.0e-1 | -2.1e-4 | ~1.0 |
| $StimulusDrive \cap Expectation$ | -5.4e-4 | 1.3e-3 | -4.4e-4 | 4.1e-3 | -9.8e-4 | 1.3e-3 | 1.4e-3 | <1.0e-5 |
| $RepetitionSuppression \cap Expectation$ | 1.1e-4 | 9.8e-2 | 1.1e-5 | 8.2e-1 | 1.2e-4 | 1.9e-1 | 4.6e-4 | 2.5e-3 |
| $StimulusDrive \cap RepetitionSuppression \cap Expectation$ | -7.3e-4 | 1.2e-3 | -9.7e-4 | 2.5e-3 | -1.7e-3 | 4.3e-3 | 1.1e-3 | 2.3e-3 |

**Table Supplement1**: **Variance partitioning results for stimulus drive, repetition suppression, and expectation models.** This table reports depth-resolved statistical results from set-theoretic variance partitioning of cross-validated model fit ($R^2$) across five auditory ROIs. All values reflect the cross-validated *unique* variance explained by each model component or the *uniquely shared* variance jointly explained by specific component combinations. Shown are mean differences across cortical depth pairs (deep vs middle, middle vs superficial, deep vs superficial) with associated paired-sample bootstrapped $p_{\text{FDR}}$ values, and one-sample bootstrap $t$-tests against zero for the ROI-level average explained variance. $p$-values for depth comparisons are FDR-corrected across depth contrasts and ROIs; ROI-level $p$-values are FDR-corrected across ROIs. Cells with $p_{\text{FDR}} > 0.05$ are shown with a grey background to denote non-significance.

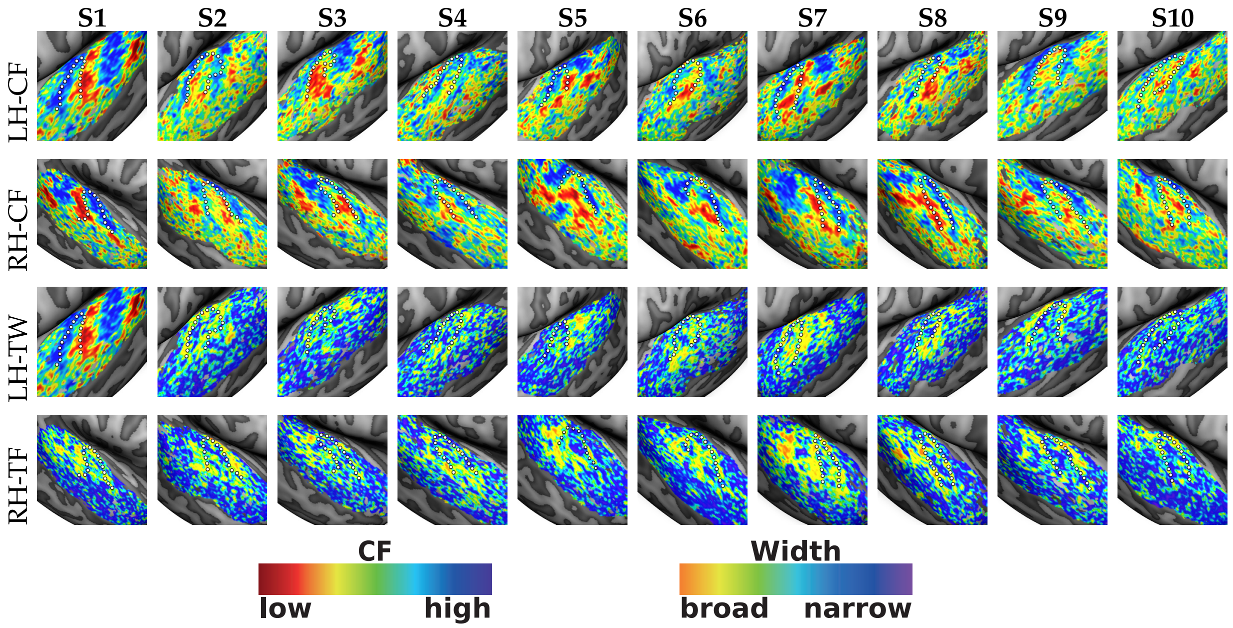


**Figure S4**: **Individual subject pRF center frequency and tuning width maps.** Rows show estimated population receptive field (pRF) maps for center frequency (CF) and tuning width (TW), separately for left (LH) and right (RH) hemispheres. Columns correspond to individual subjects (S1–S10). Maps were manually masked to auditory cortex and thresholded at $r > .2$, based on the correlation between voxelwise beta estimates and the fitted Gaussian pRF model. The dotted white line denotes the anatomical border of Heschl's gyrus.

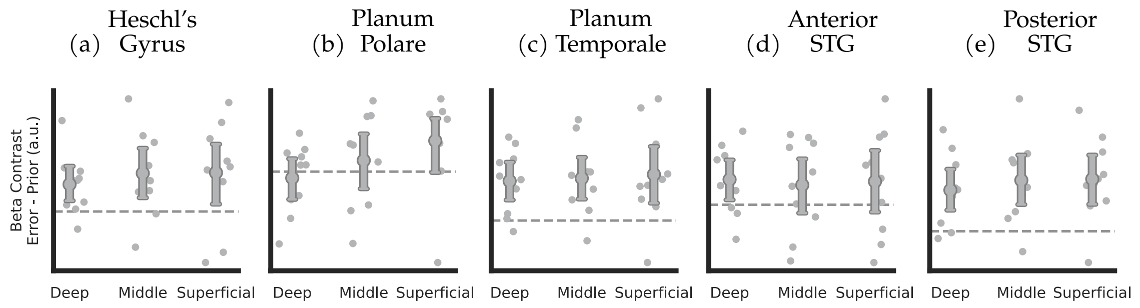


**Figure S5**: **No reliable laminar dissociation between priors and prediction errors: coefficient-level contrasts.** Each panel shows the contrast between beta coefficients for prediction errors and priors across cortical layers, assessed separately for five auditory regions of interest (ROIs): a Heschl's Gyrus (HG), b Planum Polare (PP), c Planum Temporale (PT), d anterior STG (aSTG), and e posterior STG (pSTG). This analysis tested whether coefficients related to priors and errors differed systematically in their depth profiles, using direct contrasts of their cross-validated regression coefficients. While a weak trend emerged toward larger error-related responses in superficial layers, no effects survived FDR correction ($p_{\mathrm{FDR}} > 6.8\mathrm{e}{-2}$). Importantly, these coefficients were estimated while stimulus drive and repetition suppression remained in the model, and our main analysis suggests that the shared variance between stimulus drive and expectations largely drives superficial-layer responses, which may explain the lack of unique dissociations here. Error bars indicate bootstrapped 95% confidence intervals.

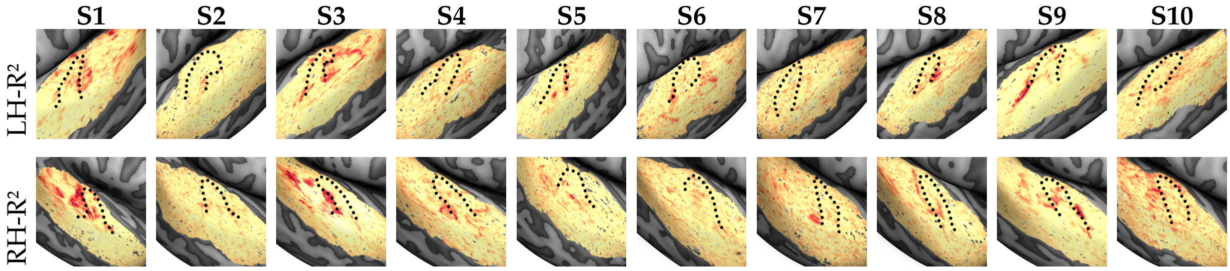


**Figure S6**: **Individual subject maps of full model fit (*Stimulus Drive* ∪ *Repetition Suppression* ∪ *Expectations*).** Shown are $R^2$ maps for each individual participant (S1–S10), separately for left (LH) and right (RH) hemispheres. The maps reflect variance explained by the full model incorporating stimulus drive, repetition suppression, and expectation components. Data were manually masked to auditory cortex and thresholded at $p < .05$ (uncorrected). The dotted black line denotes the anatomical border of Heschl's gyrus. Colour indicates model fit, with red reflecting higher $R^2$ values and yellow lower values; explained variance reaches up to approximately 12%.

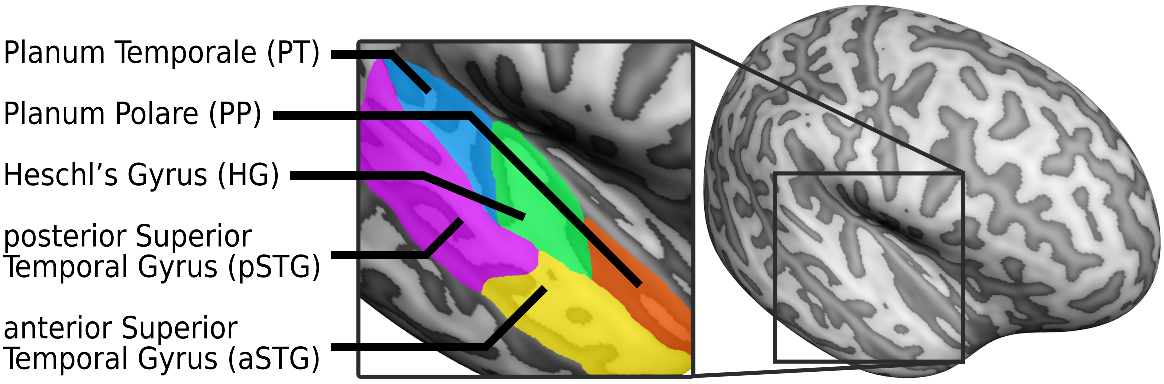


**Figure S7**: **Region of Interest (ROI) segmentation.** Lateral view of the inflated right-hemisphere mid–gray matter surface in a representative subject, with regions of interest delineated according to macroanatomical landmarks [55, 33]. The auditory cortex was subdivided into Heschl's gyrus (HG), Planum Polare (PP), Planum Temporale (PT), anterior superior temporal gyrus (aSTG), and posterior superior temporal gyrus (pSTG).

| | Deep vs Middle | | Middle vs Superficial | | Deep vs Superficial | | ROI Level | |
|---|---|---|---|---|---|---|---|---|
| | $R^2$ | $p_{\mathrm{FDR}}$ | $R^2$ | $p_{\mathrm{FDR}}$ | $R^2$ | $p_{\mathrm{FDR}}$ | $R^2$ | $p_{\mathrm{FDR}}$ |
| **Heschl's gyrus** | | | | | | | | |
| *Baseline* | -3.9e-05 | 8.8e-01 | -3.6e-04 | 1.4e-01 | -4.0e-04 | 1.4e-01 | 1.5e-03 | 2.5e-05 |
| *Priors* | 1.7e-04 | 2.2e-01 | 1.0e-04 | 4.9e-01 | 2.7e-04 | 2.3e-01 | 8.1e-05 | 2.4e-01 |
| *Errors* | 6.5e-04 | <1.0e-5 | 1.6e-04 | 4.4e-01 | 8.1e-04 | 2.3e-02 | 2.2e-04 | 6.4e-02 |
| **Planum Polare** | | | | | | | | |
| *Baseline* | -6.4e-05 | 6.4e-01 | 3.6e-05 | 8.8e-01 | -2.8e-05 | 8.9e-01 | 1.4e-03 | <1.0e-5 |
| *Priors* | -3.6e-05 | 7.0e-01 | 1.3e-04 | 2.1e-01 | 9.4e-05 | 4.9e-01 | 3.4e-04 | 9.5e-04 |
| *Errors* | 9.6e-05 | 3.6e-01 | 2.9e-04 | 6.9e-02 | 3.9e-04 | 1.3e-02 | 9.5e-04 | <1.0e-5 |
| **Planum Temporale** | | | | | | | | |
| *Baseline* | 1.3e-04 | 3.6e-01 | -3.1e-05 | 8.8e-01 | 9.9e-05 | 8.8e-01 | 1.1e-03 | 1.3e-04 |
| *Priors* | 6.2e-05 | 5.4e-01 | 2.7e-04 | 5.9e-02 | 3.3e-04 | 5.7e-03 | 6.8e-05 | 2.4e-01 |
| *Errors* | 3.4e-04 | 7.0e-03 | 4.7e-04 | 2.3e-02 | 8.1e-04 | 2.3e-02 | 3.8e-04 | 1.3e-02 |
| **anterior STG** | | | | | | | | |
| *Baseline* | 1.1e-04 | 3.8e-01 | 2.9e-04 | 1.4e-01 | 4.0e-04 | 1.4e-01 | 1.3e-03 | <1.0e-5 |
| *Priors* | 1.1e-04 | 2.3e-01 | 2.1e-04 | 1.0e-01 | 3.3e-04 | 3.6e-02 | 2.4e-04 | 4.4e-03 |
| *Errors* | 5.4e-05 | 4.6e-01 | 4.1e-04 | 6.8e-03 | 4.7e-04 | 1.5e-03 | 7.9e-04 | 2.5e-05 |
| **posterior STG** | | | | | | | | |
| *Baseline* | 1.6e-04 | 1.4e-01 | 6.9e-05 | 6.4e-01 | 2.3e-04 | 3.6e-01 | 1.2e-03 | <1.0e-5 |
| *Priors* | 1.3e-04 | 3.2e-01 | 7.8e-05 | 2.3e-01 | 2.1e-04 | 2.3e-01 | 2.0e-04 | 3.7e-02 |
| *Errors* | 3.2e-04 | 1.5e-02 | 1.9e-04 | 6.0e-02 | 5.1e-04 | 1.8e-03 | 7.2e-04 | 4.6e-03 |

**Table Supplement2**: **Variance partitioning results for baseline (stimulus drive with repetition suppression), priors, and errors.** This table reports depth-resolved statistical results from set-theoretic variance partitioning of cross-validated model fit ($R^2$) across five auditory ROIs. All values reflect the cross-validated *unique* variance explained by each model component or the *uniquely shared* variance jointly explained by specific component combinations. Shown are mean differences across cortical depth pairs (deep vs middle, middle vs superficial, deep vs superficial) with associated paired-sample bootstrapped $p_{\mathrm{FDR}}$ values, and one-sample bootstrap $t$-tests against zero for the ROI-level average explained variance. $p$-values for depth comparisons are FDR-corrected across depth contrasts and ROIs; ROI-level $p$-values are FDR-corrected across ROIs. Cells with $p_{\mathrm{FDR}} > 0.05$ are shown with a grey background to denote non-significance.
